## Supplemental Text and Figures for "Cost-effective DNA methylation profiling by FML-seq"

### Supplemental text: Prevention of adapter dimers

FML-seq uses sequencing adapters with $5'$ overhangs of 4 random bases (4N) for efficient sticky-end ligation to the corresponding overhangs of unknown bases resulting from digestion by a restriction endonuclease whose motif includes methylated cytosine. This would be expected to result in overwhelming byproduct from dimerization of adapters that ligate directly to each other without a gDNA insert. FML-seq's library protocol uses a combination of several techniques to prevent this behavior:

- The adapters are added to the gDNA before digestion. After digestion there is a heat-denaturation of the restriction endonuclease, and at this step adapters that annealed to each other during storage at higher concentration may melt apart and reanneal to the ends of new gDNA fragments instead.
- The adapters lack a phosphate at the $5'$ terminus, which is required for DNA ligase to connect that terminus to a matching $3'$ hydroxyl. Thus neither strand in an adapter dimer can be ligated, and even if two adapters with complementary overhangs temporarily anneal, 4 bp of hydrogen bonding may not keep them together when the temperature is raised in subsequent steps.
- Before PCR the same polymerase is used to fill in the second strand of the adapter by extension from the nick where the adapter's $5'$ end is not joined to the gDNA insert's $3'$ end. Even if an adapter dimer holds together at the PCR polymerase's extension temperature, it will also have a nick on the opposite strand, so the polymerase would have to jump over a gap in the template strand to fill in the new strand.
- The $5'$ end of the stem-loop (hairpin) adapter has the 4N sticky-end overhang and a complementary sequence that forms the double-stranded stem. In addition to PCR suppression this stem could also cause adapter dimerization by melting and reannealing to a different molecule. However, in the stem sequence all thymine bases are replaced with uracil, and a dU-intolerant proofreading polymerase is used to prevent the stem sequence from being replicated.
- Furthermore the stem sequence's uracils are excised before PCR by uracil-DNA glycosylase (UDG), leaving abasic sites that further deter replication and may be fully destroyed by hydrolysis at PCR denaturation temperature. In this protocol, UDG from a hyperthermophile (*Archaeglobus fulgidus*) is included in the combined ligation/loop-breaking/PCR master mix, because the traditional *Escherichia coli* UDG interferes with the low-temperature ligation and would need to be added in an additional step between ligation and PCR, whereas the hyperthermophilic UDG appears inert at ligation temperature.
- For the Illumina sequencing platform, the adapters are based on the less traditional but equally well supported Nextera sequence (Tn5 transposon) rather than standard TruSeq, because that sequence is T-rich on the $5'$ strand and therefore this protocol's adapters are U-rich in the portion removed by UDG.
- These adapter sequences are incomplete and require long PCR primers to extend them to full length with multiplexing indexes. Like the Nextera design, the PCR primers are also incomplete and only partially overlap the ligated adapter sequences. Thus both an adapter and a PCR primer are required to create a full-length product matching both the flowcell's amplification primers and the sequencing primers, so neither the original adapters nor the PCR primers can form sequenceable byproducts alone.

An additional consequence of destroying the $5'$ ends of the original adapters is that the random overhang is also destroyed: even if it contains a mismatch to the complementary gDNA sequence that is tolerated by the ligase, the mismatched base is not part of the final molecule's sequence. The final sequence is derived only from the gDNA and the random overhang does not contribute mismatches at the ends of sequence reads.

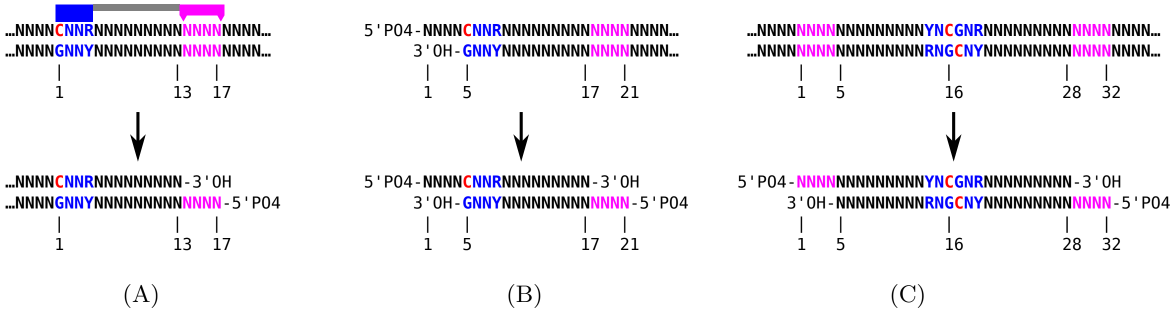


Figure S1: DNA digestion by MspJI restriction endonuclease. A: The enzyme's recognition domain (blue) binds a motif (blue) containing methylated cytosine (red), $^{m}$CNNR, while its endonuclease domain (magenta) cuts at distances of 13 and 17 bp from the methylated cytosine position. Digestion leaves a 5′ overhang of 4 nt with a terminal phosphate, and terminal hydroxyl on the 3′ underhang. B: Digestion by MspJI cannot result in very short fragments. The shortest theoretically possible library insert would be generated by recognition of a $^{m}$CNNR site immediately flanking the 4 nt overhang resulting from a previous digestion, leaving a total insert of 21 bp after ligating the adapters. In reality, MspJI may not necessarily be able to bind its motif without additional paired bases on both sides. Empirically we observe, in high-input libraries without size selection, only about 0.04% of library inserts shorter than 22 bp, 0.05% shorter than 25 bp, 0.33% shorter than 30 bp (Figure S7A). C: In the special case of a fully methylated CpG within a YNCGNR palindrome, two MspJI enzymes may digest the DNA symmetrically and produce a 32 bp insert. In FML-seq libraries from human gDNA, inserts of this length are disproportionately common but not the majority, about 4% of all inserts (Figure S7A).

### Supplemental text: Comparison with other methods based on methylation-specific digestion

Previous methods that used MspJI or another methylation-specific restriction endonuclease focused exclusively on the 32 bp fragment produced by two complete, symmetric digestions of a fully methylated CpG (Figure S1C). This requires a precise size selection such as cutting a band out of an acrylamide electrophoresis gel, and discards about 96% of the library product (see Figure S7). In contrast, FML-seq treats every cut site as informative of a methylated base, even where digestion is incomplete or the sequence motif is not palindromic.

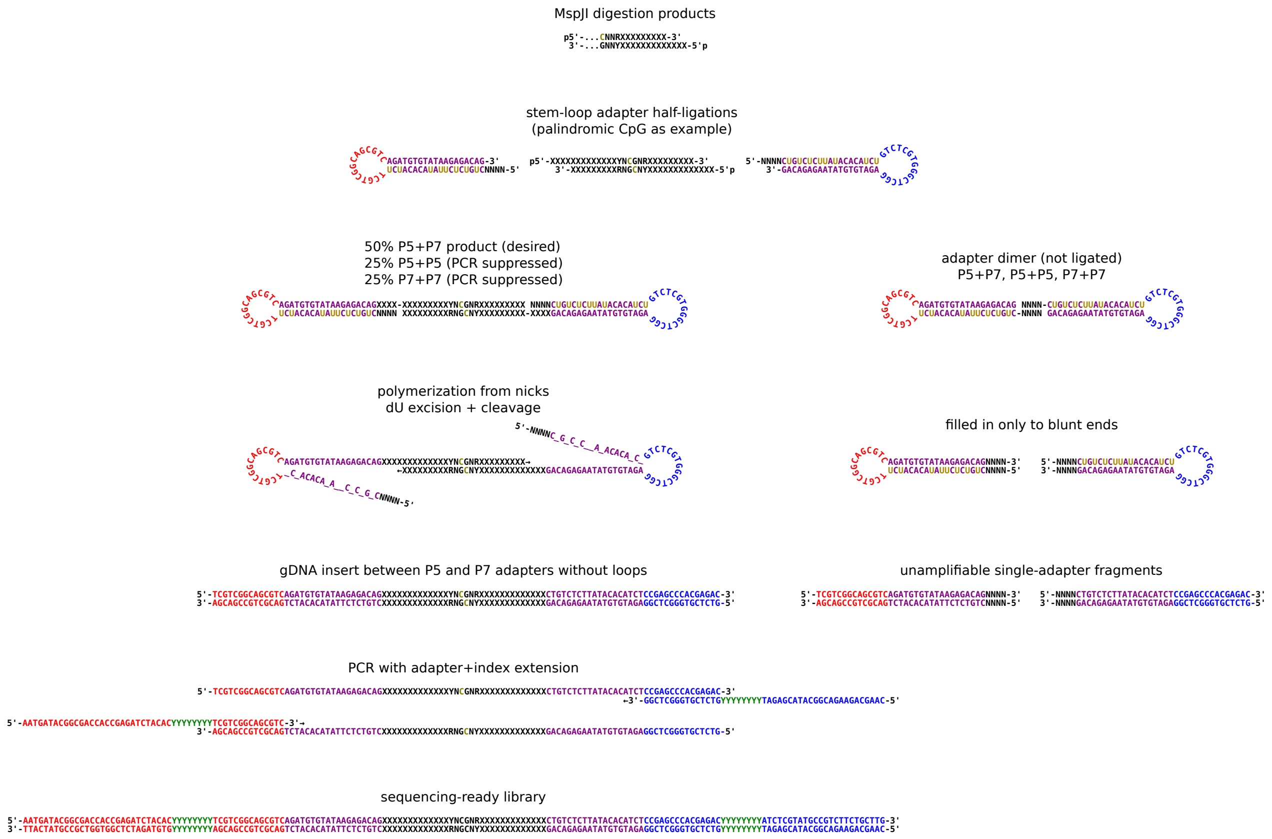


Figure S2: FML-seq molecule sequences using MspJI restriction endonuclease and Nextera sequencing adapters. For convenience a palindromic YNCGNR site with two complementary CGNR motif sites is shown, though fragments can be produced by a non-palindromic single site and a given fragment does not necessarily contain the motif sites responsible for its digestion. The stem-loop (hairpin) adapters are an equimolar mix of two versions, one for each end of a sequencing-ready library molecule (Illumina P5 and P7), but except for their slightly different sequences they are used identically and the ligation product has no polarity. By chance, half of the digestion products will be ligated between two of the same adapter (P5 & P5 or P7 & P7), producing an unsequenceable molecule that is unlikely to amplify because of PCR suppression; this limits the efficiency of the library synthesis to 50%. Nick extension replaces the second adapter strand while UDG excises uracil and the abasic sites are hydrolyzed, leaving an amplifiable library molecule. Because the stem-loop adapters lack 5′ phosphate, they ligate to a gDNA fragment on only one strand, and if two adapters anneal without a gDNA insert, the nick extension stops at the nick on the opposite strand. *Oligonucleotide sequences © 2021 Illumina, Inc. All rights reserved. Derivative works created by Illumina customers are authorized for use with Illumina instruments and products only. All other uses are strictly prohibited.*

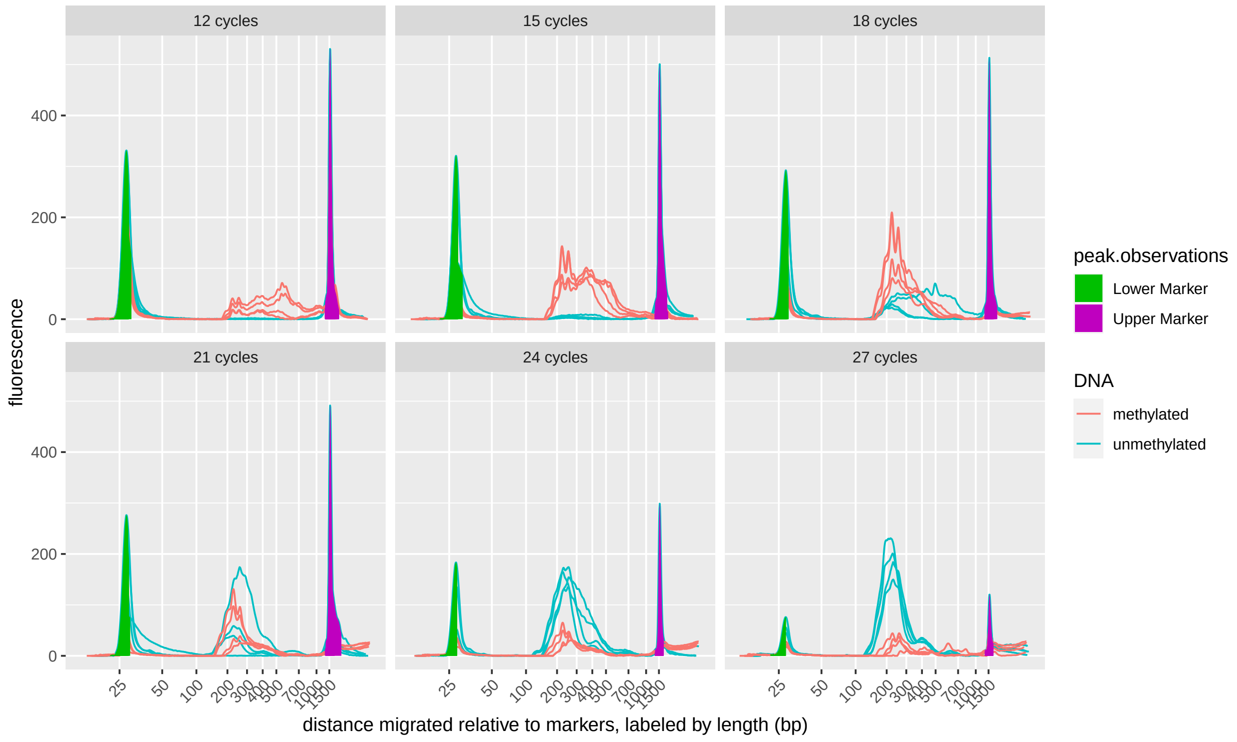


(A)

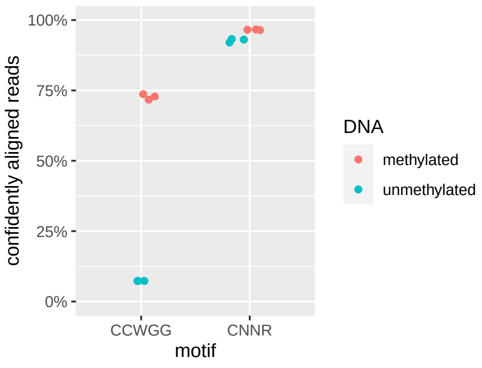


(B)

Figure S3: Specificity of FML-seq for methylated DNA. A: Electropherograms of FML-seq libraries prepared from 60 ng methylated or unmethylated lambda bacteriophage genomic DNA with varying numbers of PCR cycles, four replicates per condition. The optimized protocol uses 15 cycles for this amount of methylated gDNA, but unmethylated DNA requires about 24 cycles to produce a similar yield. Note: With more than 15 PCR cycles, the overamplified libraries from methylated gDNA are converted into slow-migrating artifacts that appear to the right of the upper marker, giving the appearance of low yield in the measurable length range. B: Alignment of sequence reads at the expected distance from motif sites. Methylation by the host's Dcm is expected at the C$^{m}$CWGG motif; however, MspJI cuts at any $^{m}$CNNR.

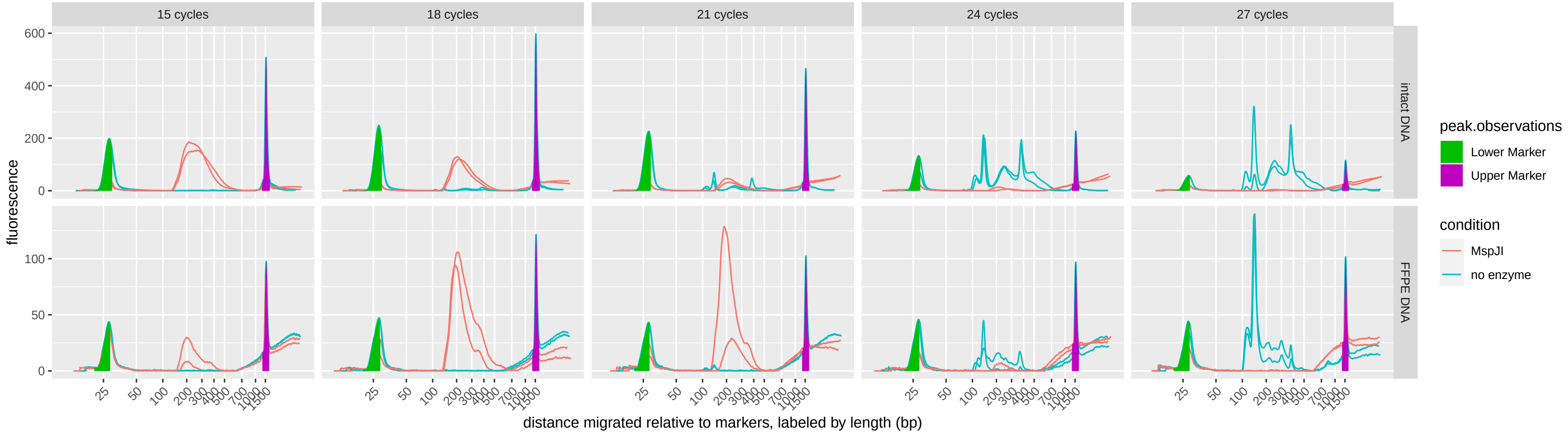


Figure S4: Specificity of FML-seq for digested gDNA: Electropherograms of FML-seq libraries prepared from 60 ng intact human gDNA from fresh cells or degraded human gDNA from formalin-fixed, paraffin-embedded (FFPE) tissue, using either methylation-specific digestion by MspJI restriction endonuclease or a no-endonuclease control, two replicates per condition. The no-endonuclease control shows no library yield unless the number of PCR cycles is greatly increased, at which point its yield is similar to a no-DNA control (compare Figure S5C). Note: As in Figure S3A, the yields of libraries from the MspJI condition appear to decrease with high PCR cycles due to overamplification artifacts; furthermore, undigested gDNA fragments from the FFPE sample are visible near and beyond the upper marker because of their short length, even where no library is detectable.

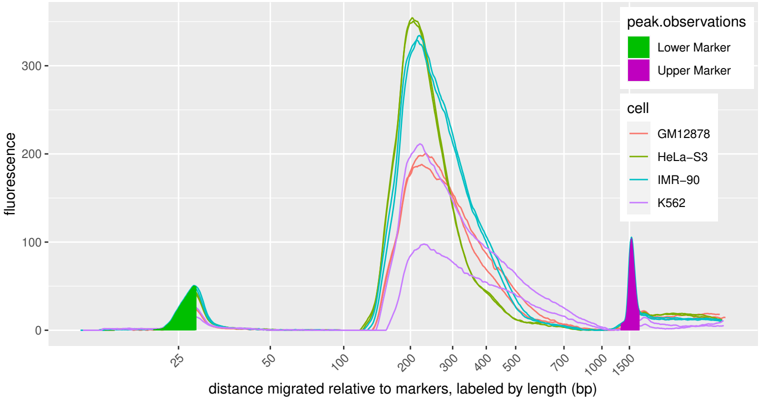


(A)

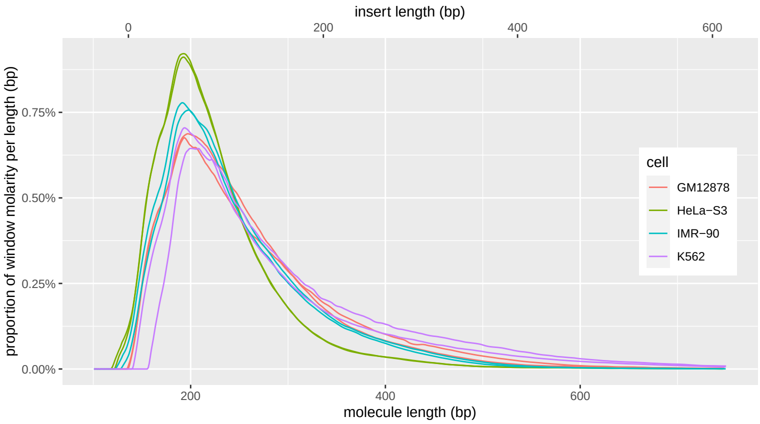


(B)

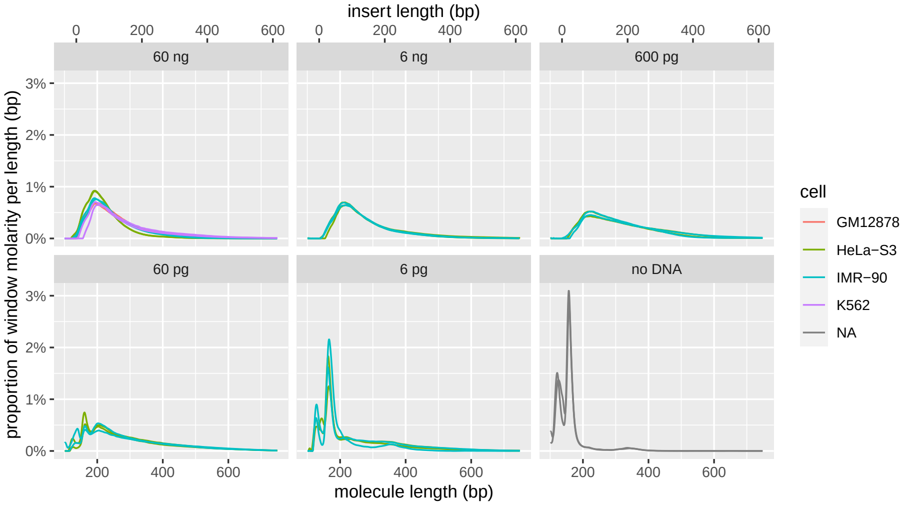


(C)

Figure S5: Properties of FML-seq libraries from human cell-line gDNAs. A: Electropherograms of libraries from 60 ng gDNA, two replicates per cell line. B: Molar length distributions on a linear axis, normalized to equal totals within the graphing window. Insert length is calculated by subtracting the combined length of the indexed sequencing adapters, 136 bp, from the molecule length. The TapeStation lacks sufficient resolution to show the expected peak of 32 bp inserts (168 bp molecules). C: Libraries from serial-dilution gDNA samples and no-DNA control.

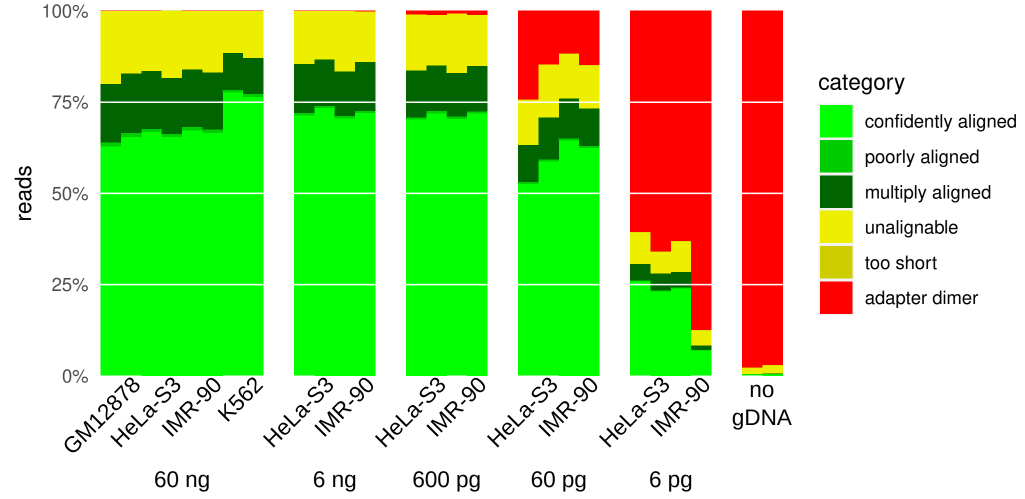


(A)

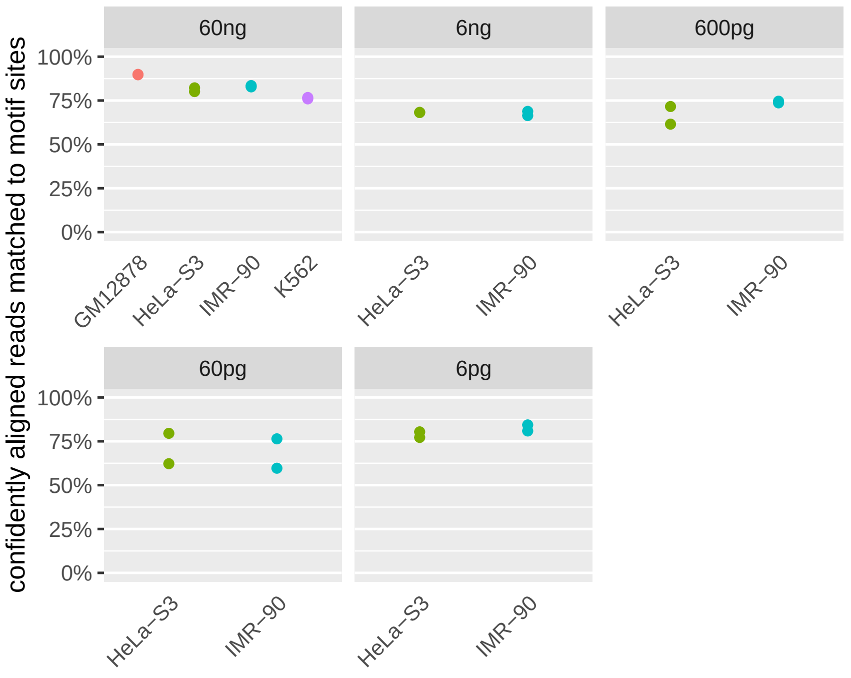


(B)

Figure S6: Properties of FML-seq sequence reads from human cell-line gDNAs. A: Alignability of read sequences to the reference genome. “Adapter dimers” are inserts of 10 bp or less and “too short” reads are longer than adapter dimers but shorter than `bwa-mem2`’s minimum seed length of 19. “Confidently” and “poorly aligned” are uniquely aligned reads above or below the minimum MAPQ of 10 (posterior probability of correct alignment 0.9). B: Alignment of sequence reads at the expected distance from CGNR motif sites.

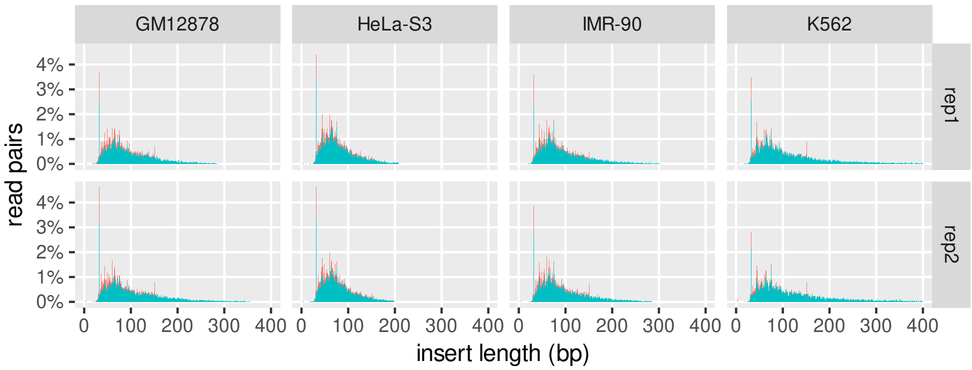


(A)

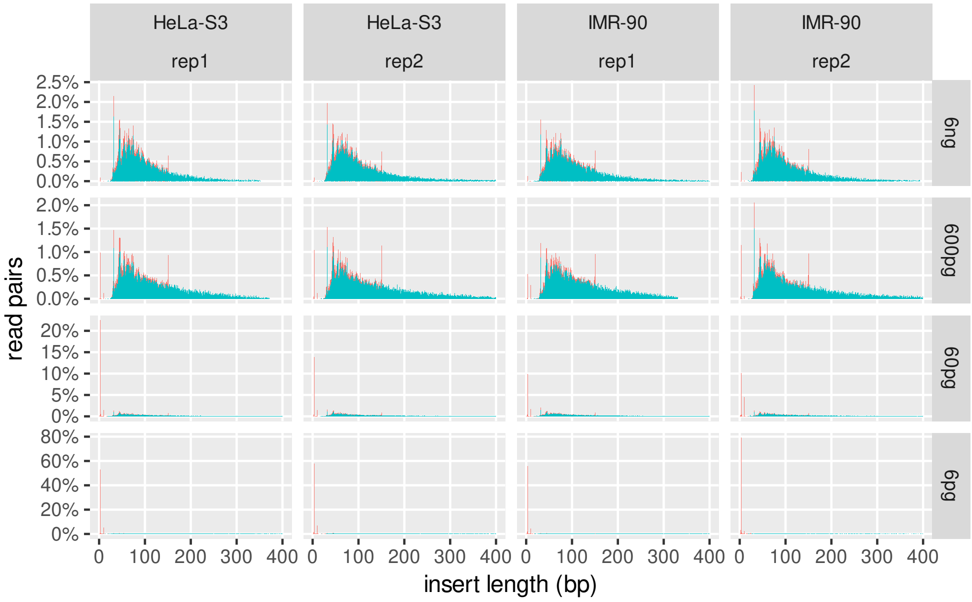


(B)

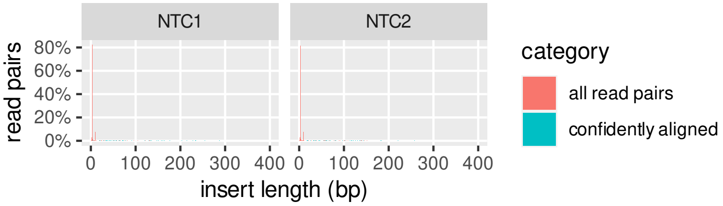


(C)

Figure S7: Lengths of FML-seq library inserts inferred from sequencing. Lengths of ”all read pairs” are inferred by adapter trimming, but this is limited to inserts shorter than the sequence read length, 154 nt (untrimmed reads are not shown). Lengths of ”confidently aligned” read pairs (MAPQ $\geq$ 10) are inferred by aligned positions in the reference genome. For direct comparison the denominator of both fractions is the total number of post-filter sequenced read pairs, i.e. ”all read pairs” sum to 100% but ”confidently aligned” read pairs sum to the proportion of confidently alignable read pairs in each library, and for any given insert length below the read length the relative height of the ”confidently aligned” bar reflects the proportion of ”all read pairs” at that length. There are prominent peaks at 4 bp from adapter dimers (no insert of gDNA sequence); at 32 bp from symmetrically digested, fully methylated CpG sites; and at 151 bp due to the 3-base minimum for detection of adapter sequence. A: All tested cell lines, 60 ng gDNA each. B: Serial dilutions of HeLa-S3 and IMR-90. C: No-template controls (no gDNA).

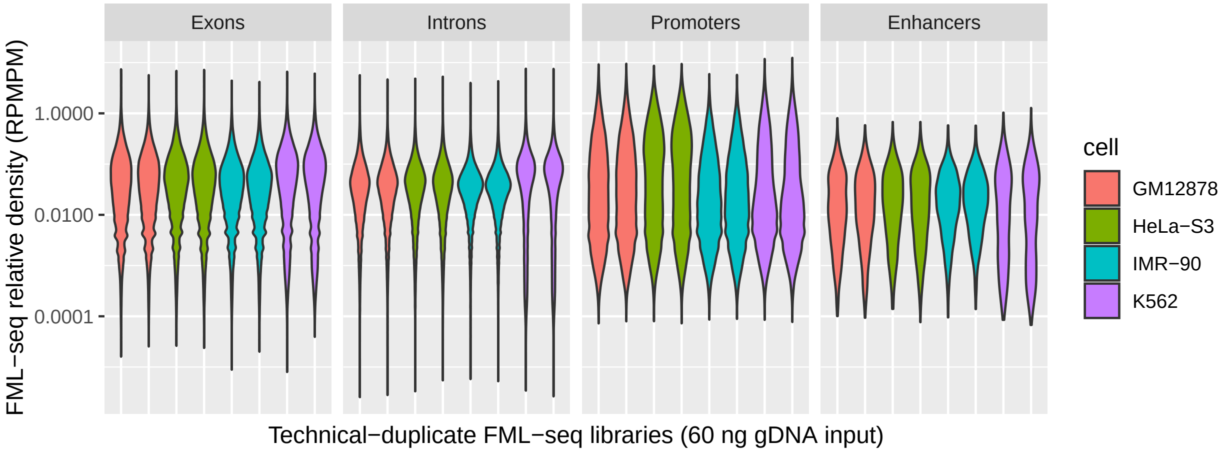


(A)

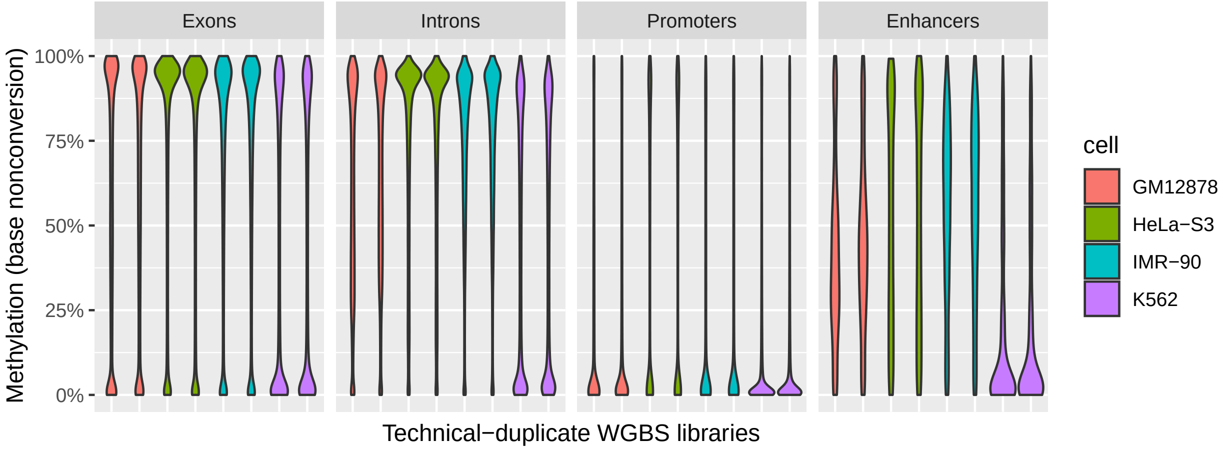


(B)

Figure S8: Distribution of methylation scores among different categories of annotated functional genome elements. A: Density of sequence reads from FML-seq assigned to CGNR sites within the genome elements. Here reads per million per motif (RPMPM) is normalized against the total number of reads assigned to all CGNR sites, rather than the sum within only a given category of genome elements, in order to compare the different categories despite covering widely different proportions of the genome (286,435 exons, 297,593 introns, 40,351 promoters, 1,402 enhancers). B: Methylation of CpG sites within the genome elements according to whole-genome bisulfite sequencing.

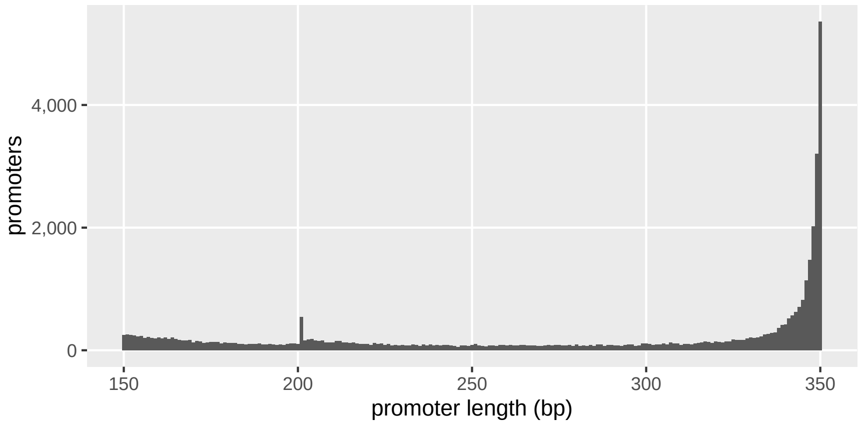


(A)

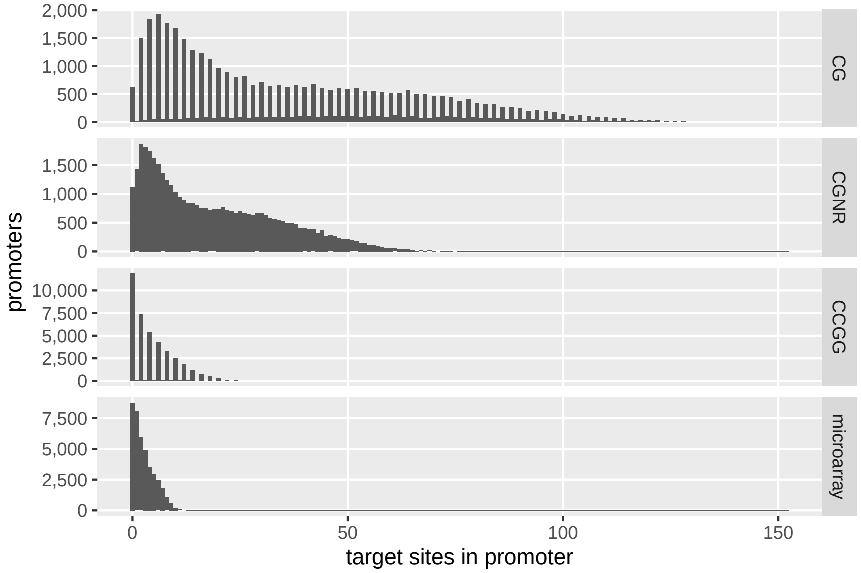


(B)

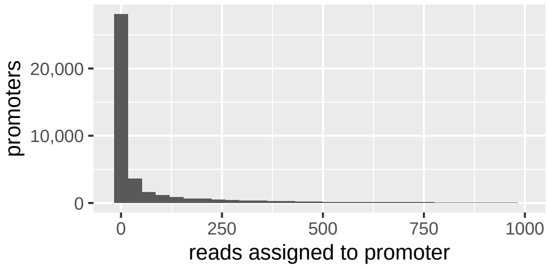


(C)

Figure S9: Properties of the 40,351 annotated human promoters used for data analysis. A: Lengths of promoter regions in the reference genome. B: Number of targets for each DNA methylation profiling method per promoter (CG: canonical human DNA methylation; CGNR: the MspJI restriction endonuclease used to target methylated cytosine in FML-seq; CCGG: the MspI restriction endonuclease used to reduce the number of targets in RRBS). Both copies of palindromic motifs CG and CCGG are counted unless the potentially methylated CpG straddles the boundary of the promoter, resulting in counts that are nearly always even. The promoters are enriched for CpG dinucleotides (average 6.5 per 100 bp) relative to the entire reference genome (1.1 per 100 bp). C: FML-seq read counts per promoter in a representative sample, K562 replicate 1. The 1,126 promoters without CGNR motif sites are included as zero counts.

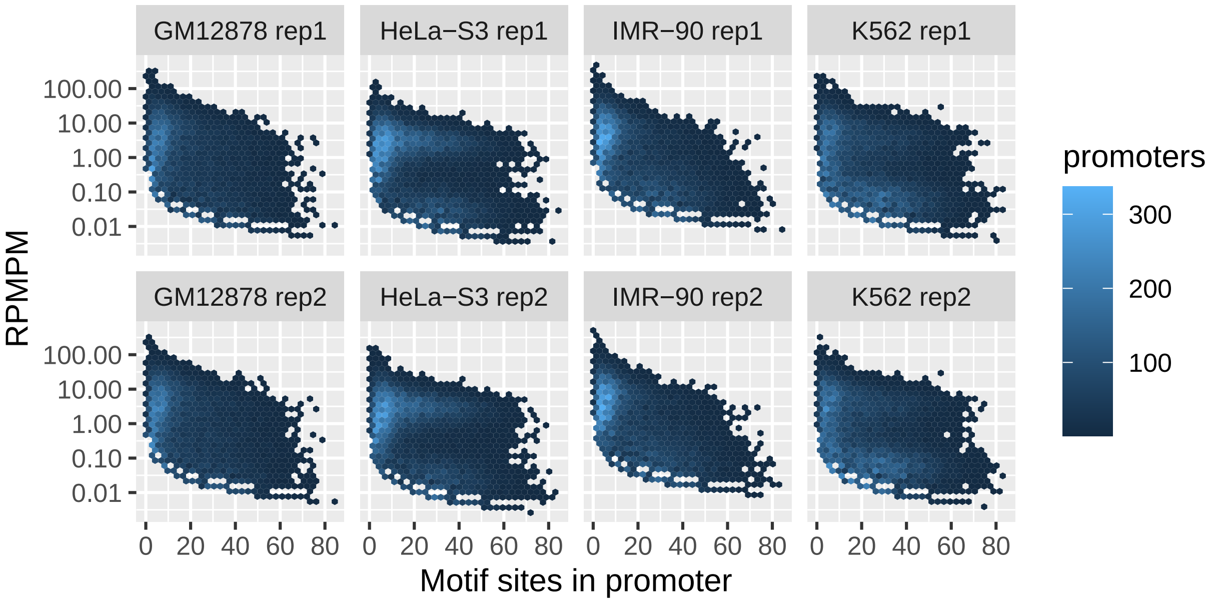


Figure S10: FML-seq reads per million per motif (RPMPM) by number of restriction motif sites in the promoter.

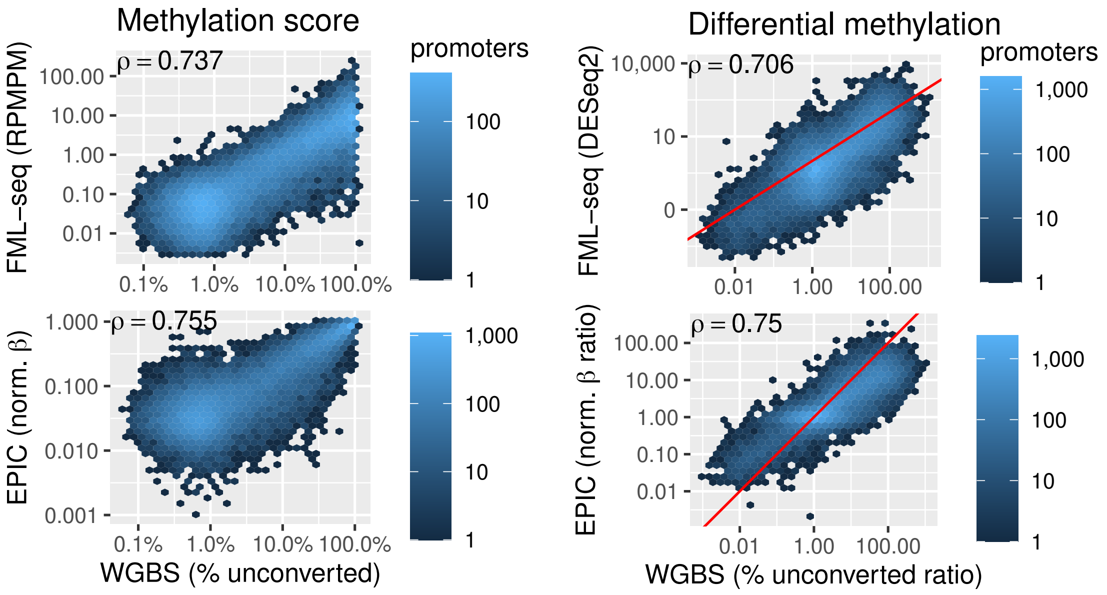


(A)

(B)

Figure S11: Different platforms' methylation scores in K562 (A) and fold changes for K562 vs. HeLa-S3 (B), at all promoters. The red line shows $y = x$.

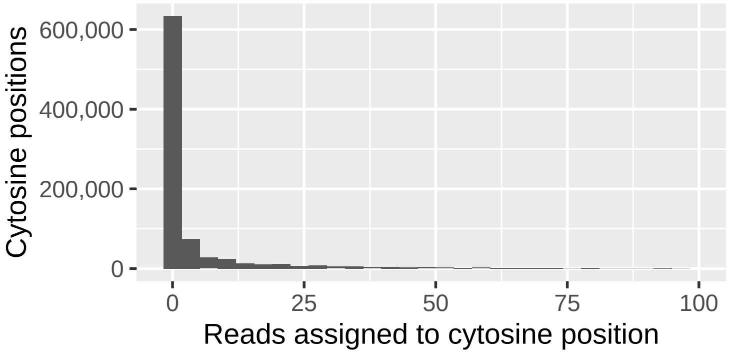


(A)

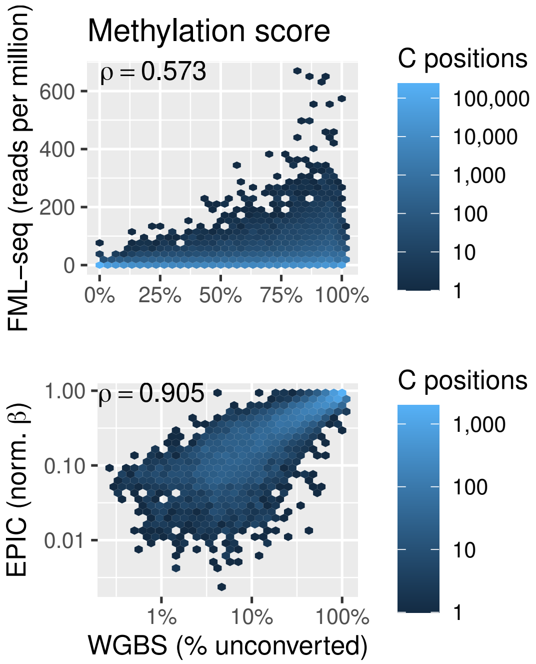


(B)

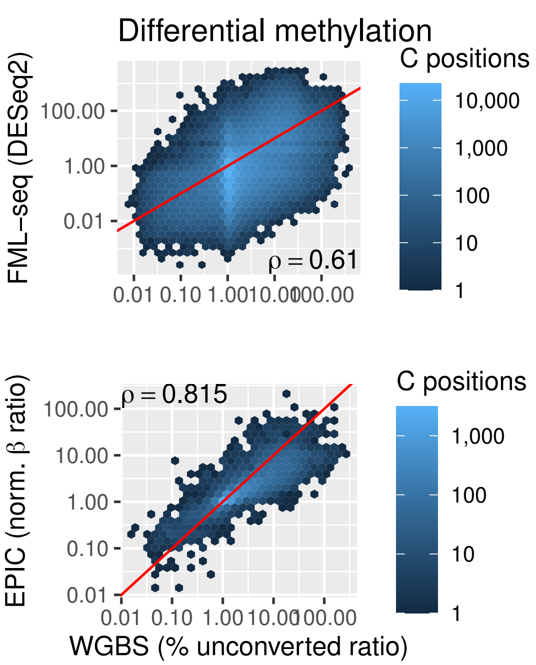


(C)

Figure S12: Analysis of FML-seq at individual cytosine positions. Only cytosines within CpG dinucleotides are considered overall and only the first cytosine of the CGNR motif is considered for FML-seq read counts. Only chromosome 22, the shortest autosome, is considered in order to reduce the scale of the analysis (1,686,636 cytosine positions, 874,441 in CGNR). A: FML-seq read counts per cytosine position in a representative sample, K562 replicate 1. A: Different platforms' methylation scores in K562. B: Different platforms' fold changes for K562 vs. HeLa-S3, at all cytosine positions with CpG dinucleotides on chromosome 22, the shortest autosome. The red line shows $y = x$.

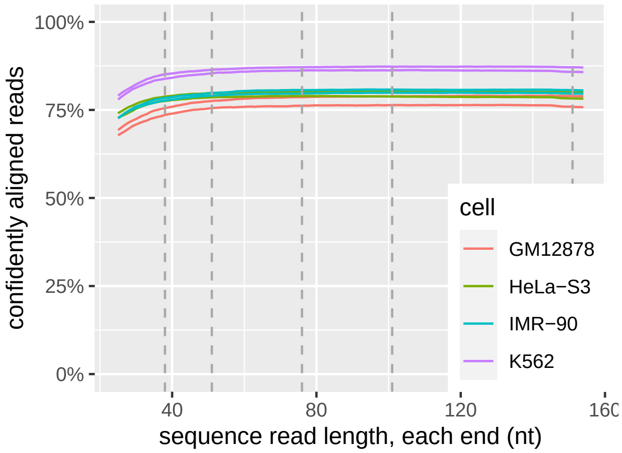


Figure S13: Long sequencing reads are unnecessary for FML-seq. Reads shown are from the MiSeq v2 sequencing kit at 2×154 nt. Confident alignment is MAPQ $\geq$ 10. Commonly available read lengths are marked: 2×38, 2×51, 2×76, 2×101, 2×151.

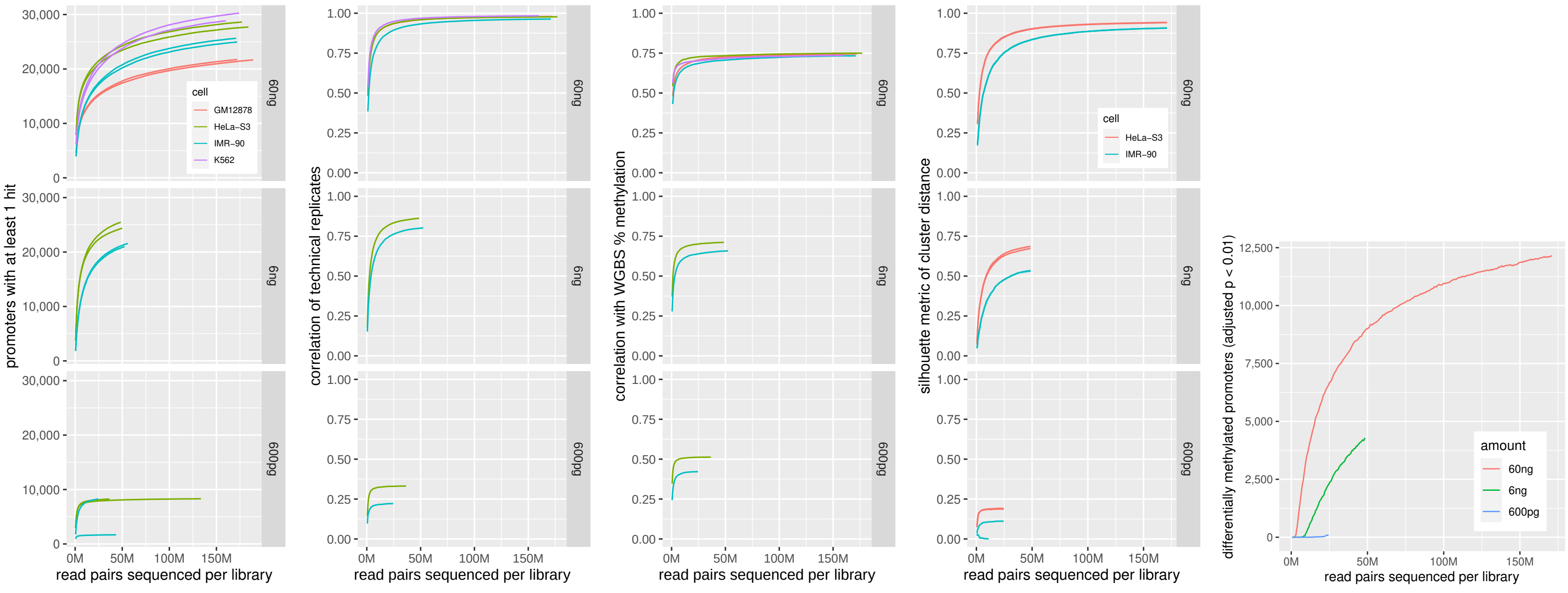


(A) (B) (C) (D) (E)

Figure S14: Deep sequencing is unnecessary for FML-seq. Lower sequencing depths were simulated by subsampling the full data. A: Promoters with at least 1 FML-seq read attributed to a CGNR motif site within the promoter boundaries. B: Pearson correlation of log RPMPM across all promoters. C: Spearman correlation of pooled log RPMPM with WGBS pooled percent methylation from the same cell line across all promoters. D: Silhouette metric of cluster distance (Pearson distance $1 - r$ to the nearest member of another class $\div$ mean distance within the same class), considering only the two cell types used in serial dilutions, HeLa-S3 and IMR-90, for fair comparison. E: Promoters with significantly differential methylation (DESeq2 BH-adjusted $p < 0.01$) between HeLa-S3 and IMR-90. The number of promoters with nonzero read counts or statistical significance inevitably increases with total sequencing depth as smaller methylation signals and effect sizes become detectable; however, replicate correlations and silhouettes indicate the total amount of usable information in the sequencing library is reached more quickly as additional depth yields only duplicate reads.
