## Supplemental Protocol for "Cost-effective DNA methylation profiling by FML-seq"

xml version="1.0" encoding="UTF-8"?

**FML-seq protocol**  
Joseph W. Foley  
12 Jan 2023

- Introduction
  - Abstract
  - Experimental design considerations
  - Choosing the number of PCR cycles
- Advance preparation
  - Required equipment
  - Consumables
  - Reagents
  - Oligonucleotide dilutions
  - Reagent premixes
  - Thermal cycler programs
    - Program 1, Digestion
    - Program 2, Synthesis
  - Sample preparation
- Procedure
  - Digestion
  - Library synthesis
  - Cleanup
- Next steps
  - Quantification and validation of the libraries
  - Sequencing
  - Data processing
- Oligonucleotide designs (Illumina-specific)
  - Abbreviations
  - Library synthesis adapters
  - Indexing PCR primers
- Quick reference protocol
  - Digestion
  - Synthesis
  - Cleanup

---

### Introduction

#### Abstract

Fragmentation at methylated loci and sequencing (FML-seq) generates a sequencing library,
from pre-isolated genomic DNA, in which every sequenceable molecule indicates two
methylated cytosine positions in the genome. The FML-seq signal at any given site
in the genome is proportional to the fraction of genome copies that were methylated
at that site in the sample. Unlike other protocols for DNA methylation profiling based
on cytosine deamination by bisulfite or APOBEC, FML-seq does not require harsh chemical
conditions or numerous enzymatic reactions, nor does it require an initial DNA fragmentation
by sonication. Instead there are only three hands-on steps: digestion of the gDNA
by a methylation-specific restriction endonuclease, single-step library synthesis,
and cleanup. The entire protocol from gDNA to a sequencing-ready library can be completed
in two hours and the reagent costs are minimal.

#### Experimental design considerations

FML-seq is intended for experiments with statistically powerful numbers of replicates
per condition. Single-sample volumes in this protocol are impractical to pipet accurately
and are intended to be prepared in a master mix with excess volume. In large experiments
it may be useful to aliquot the master mix into a strip of tubes for multichannel
pipetting, though this requires a greater excess volume. When using a multichannel
pipet to dispense the reagents, consider randomizing or blocking the positions of
samples in your plate to avoid confounding any experimental variable with channels
of the pipet.

This method has been biologically validated with 6 ng to 60 ng of input gDNA (1,000
to 10,000 human cells). Lower amounts generate seemingly high-quality libraries but
they do not contain enough distinct molecules for useful data. Higher amounts have
not been tested. To prevent bias by the amount of input, dilute all gDNA samples to
the lowest concentration in the experiment and use the same amount of gDNA for all.
To normalize the amounts accurately, you must quantify the gDNA by fluorometry (Qubit,
RiboGreen) instead of spectrophotometry (NanoDrop UV-Vis).

#### Choosing the number of PCR cycles

Compute the number of required PCR cycles based on the amount of genomic DNA input,
with the formula

> Cycles = 21 − log(ng genomic DNA) ÷ log(1.9)

###### PCR cycle calculator

ng genomic DNA per sample  
Recommended:  cycles


---

### Advance preparation

#### Required equipment

- Adjustable-volume micropipets
- Programmable thermal cycler(s) for 0.2 mL PCR tubes or microplates
- Vortex mixer(s) for various tube sizes, e.g. Fisher #14-955-151
- Mini-centrifuge(s) for PCR tubes or microplates (depending on experiment scale), e.g.
  Ohaus #FC5306, Fisher #14-100-143
- Magnetic separation block for 0.2 mL PCR tubes or microplates, e.g. V&P #772F4-1,
  #771LD-4CS
- Recommended: Electronic multichannel micropipets, e.g. Eppendorf Xplorer
- Recommended: Separate workstations for pre-PCR and post-PCR steps, to avoid contamination
- Recommended: Head-mount magnifier (loupe) for easier sight of microliter volumes

Note: Especially when working with large batches in microplates, multichannel pipets
reduce handling time, hand strain, and the risk of error. Electronic pipets are especially
useful both for mixing liquids by pipetting and for quickly dispensing aliquots of
master mix. Most of the protocol involves volumes of 10 µL or less except the final
cleanup, so the Eppendorf Xplorer 0.5–10 µL and 5–100 µL models are sufficient, but
a 200 or 300 µL multichannel pipet for the final ethanol wash can be manual rather
than electronic.

#### Consumables

- 0.2 mL PCR tubes or microplates, low-retention recommended, e.g. Axygen #PCR-02-L-C,
  Eppendorf #0030129504 (semi-skirted format may be required for compatibility with
  thermal cyclers)
- Required only for microplates: PCR plate seals, e.g. Bio-Rad #MSB1001
- Disposable filtered pipet tips, low-retention strongly recommended except where indicated,
  USA Scientific TipOne RPT Filter Tips (do not confuse with regular TipOne or unfiltered),
  Eppendorf epTIPS LoRetention (do not confuse with regular epTIPS).
- Optional: 0.2 mL PCR tube strips, to hold master mix aliquots for multichannel pipetting,
  e.g. USA Scientific #1402-4780

Note: There are few choices available for low-retention PCR tubes and microplates;
we have made successful libraries in lower-quality vessels, such as 8-tube strips.
Many brands of pipet tips are claimed to be low-retention but in our experience some,
e.g. Biotix, are still so retentive that they visibly fail to aspirate small volumes
accurately. We use the "10 µL XL graduated" TipOne RPT filter tips with our 2.5, 10,
and 20 µL Eppendorf pipets and "200 µL graduated" with 100 and 200 µL Eppendorf pipets,
which take "universal"-shape tips; for Rainin-shape tips, we have heard but not validated
a recommendation for Thomas Scientific SHARP tips. Almost all low-retention tips used
in this protocol are 10 µL size except the 100/200 µL tips used briefly for mixing
by pipetting in the cleanup step, when low retention may not be as crucial. The 200/300
tips used for the final ethanol wash and 1000 µL tips used for preparing premixes
do not need be low-retention.

#### Reagents

All chemicals must be molecular biology-grade (certified free of nucleic acids and
nucleases) except where otherwise specified. Some products are available with better
pricing on larger scales than the given catalog numbers.

- **MspJI restriction enzyme**, 5 U/µL, NEB #R0661S (includes rCutSmart Buffer and Enzyme Activator)
- **T4 DNA ligase**, 5 Weiss U/µL, e.g. Thermo Fisher #EL0011
- ***Archaeoglobus fulgidus* uracil-DNA glycoylase (Afu UDG)**, 2 U/µL, NEB #M0279S (do not confuse with standard *E. coli* UDG; NEB discontinued the large scale of this product but FML-seq requires a large
  amount, so ask for a volume discount); includes ThermoPol II Reaction Buffer, which
  is not used in this protocol
- **Kapa HiFi HotStart polymerase plus dNTPs**, Roche #07958889001 (do not confuse with 2X ReadyMix or with Uracil+ enzyme); includes
  enzyme, Fidelity Buffer, dNTP mix, and other reagents not used in this protocol
- **SPRI bead mix**, Beckman Coulter #A63880 or equivalent or homemade (note: SPRI is not used for size selection so more expensive bead mixes validated
  for size selection such as SPRIselect are not required; however, some bead mixes from
  other vendors are more dilute and require a different volume ratio)
- **P5 and P7 adapters and indexed thiolated PCR primers** ordered as custom oligonucleotides (see "Oligonucleotide designs")
- **Molecular biology-grade water**, nuclease-free, CAS #7732-18-5, e.g. Thermo Fisher #AM9932
- **Reagent-grade ethanol and ultrapure water** (e.g. Milli-Q) for only the 80% ethanol wash, to conserve certified molecular biology-grade
  water, but they must be freshly mixed
- **ATP solution, Tris-buffered**, 100 mM, e.g. Thermo Fisher #R1441 (do not store unbuffered NTPs)
- **Disodium EDTA (Na_2_EDTA)**, 500 mM solution, CAS #139-33-3, e.g. Thermo Fisher #R1021
- **Polyethylene glycol, molecular weight 4000 to 8000**, included in Thermo Fisher #EL0011 or e.g. MilliporeSigma #P1458
- **DNA storage buffer**, e.g. 1X TE + 0.05% Tween 20

#### Oligonucleotide dilutions

Resuspend lyophilized oligonucleotides in DNA storage buffer at about 100 μM. Optionally
verify by UV spectrophotometry (measure 1/10 dilutions for better accuracy) that the
yields are within 10% of expectations.

#### Reagent premixes

To simplify the routine protocol, premix some of the reagents in large batches. Make
sure to fully thaw, vortex, and (if possible) briefly centrifuge all reagents before
mixing them, then vortex again to mix and centrifuge the final solution. Store the
mixes at –20 °C.

###### Oligo Mix

samples: 

|  |  |
| --- | --- |
| DNA storage buffer | µL |
| NEB Enzyme Activator, 15 µM | µL |
| FML-seq P5 adapter, 100 µM | µL |
| FML-seq P7 adapter, 100 µM | µL |

total:  µL

Adjust the volumes of adapter stocks used here according to their measured concentrations
(see Oligonucleotide dilutions).

###### Synthesis Buffer

samples: 

|  |  |
| --- | --- |
| Kapa Fidelity Buffer, 5X | µL |
| water | µL |
| disodium EDTA, 500 mM | µL |
| Kapa dNTP mix, 10 mM each | µL |
| ATP, 100 mM | µL |
| PEG 4000, 50% m/v | µL |

total:  µL

PEG solution is viscous; slowly reverse-pipet it.

###### Synthesis Enzymes

samples: 

|  |  |
| --- | --- |
| Afu UDG, 2 U/µL | µL |
| Kapa HiFi HotStart polymerase, 1 U/µL | µL |
| T4 ligase, 5 Weiss U/µL | µL |

total:  µL

**PCR primers**: dilute each pair together to **3 µM each primer** in TE+Tween.

#### Thermal cycler programs

Note: The final holds are safe stopping points and the reaction products are stable,
so you can choose a final holding temperature to minimize the instrument's energy
consumption and fan noise. For example, 14 °C is the closest to room temperature at
which an Applied Biosystems Veriti disables its heated lid. If you are nearby when
the program finishes, you can transfer the samples to a refrigerator in order to power
off the thermal cycler until you are ready for the next step.

##### Program 1, Digestion

- 37 °C hold (pre-heat, then place samples in thermal cycler and end hold)
- 37 °C 30:00
- 65 °C 5:00
- 10–25 °C hold

Reaction volume: 5 µL

##### Program 2, Synthesis

- 16 °C hold (pre-cool, then place samples in thermal cycler and end hold)
- 16 °C 5:00
- 72 °C 3:00
- 98 °C 0:45
- cycles:
  - 98 °C 0:15
  - 60 °C 0:30
  - 72 °C 0:15
- 72 °C 1:00
- 10–25 °C hold

Reaction volume: 10 µL

The number of PCR cycles is determined by the formula and may vary from one experiment to another.

#### Sample preparation

Genomic DNA must be free of histones and other proteins, e.g. isolated by a protocol
that includes proteinase K treatment. Preferably it should not be incubated above
70 °C, to keep it double-stranded and avoid GC-content bias. DNA is best stored in
a slightly basic buffer and a surfactant improves pipetting accuracy (see "Reagents"). This protocol generates successful libraries from formalin-fixed, paraffin-embedded
(FFPE) samples but has not been biologically validated for that application.

---

### Procedure

#### Digestion

Decontaminate the pre-PCR workstation.

Thaw the gDNA samples and fragmentation and digestion reagents to room temperature.

Pre-warm the pre-PCR thermal cycler with **Program 1, Digestion**.

Vortex and centrifuge all reagents and samples.

Combine the ingredients for Digestion Mix, then vortex and centrifuge it:

###### Digestion Mix

reactions: 

|  |  |
| --- | --- |
| water | µL |
| rCutSmart Buffer, 10X | µL |
| premade Oligo Mix | µL |
| MspJI, 5 U/µL | µL |

total:  µL

Note: The water volume may be reduced to use a greater volume of gDNA; change the
aliquot volumes accordingly.

Aliquot **4 µL Digestion Mix** to a separate tube for each sample.

Add **1 µL gDNA sample** to each tube.

Mix each sample by pipetting **3–4 µL up and down 5 times**.

Seal and centrifuge the samples.

Place the samples in the thermal cycler and end the 37 °C hold.

Return the digestion reagents to the freezer.

Safe stopping point: After digestion, the samples can stay in the thermal cycler or
a refrigerator for a few days before you continue to library synthesis.

#### Library synthesis

Thaw the library synthesis reagents to room temperature.

Pre-cool the PCR thermal cycler with **Program 2, Synthesis** using the appropriate number of cycles.

Combine the ingredients for Synthesis Mix, then vortex and centrifuge it:

###### Synthesis Mix

reactions: 

|  |  |
| --- | --- |
| premade Synthesis Buffer | µL |
| premade Synthesis Enzymes | µL |

total:  µL

When Program 1 has reached the final hold, remove and centrifuge the samples.

Add **4 µL Synthesis** to each sample tube.

Add a different **1 µL indexed PCR primer pair, 3 µM each primer** to each sample tube.

Mix each sample by pipetting **6–8 µL up and down 5 times**.

Seal and centrifuge the samples.

Place the samples in the PCR thermal cycler and end the 16 °C hold.

Return the PCR reagents to the freezer.

Safe stopping point: After digestion, the samples can stay in the thermal cycler or
a refrigerator for a few days before you continue to cleanup.

#### Cleanup

Thoroughly resuspend the SPRI bead mix by vortexing.

When Program 2 has reached the final hold, remove and centrifuge the samples.

Add **18 µL SPRI bead mix** to each sample and **mix by pipetting** the same volume up and down 5 times.

Incubate **1 min**, then briefly centrifuge the samples to eliminate bubbles.

Place the samples on the magnet to separate, which may take several minutes. A white
paper towel under the magnet makes it easier to see the bead pellets.

Prepare fresh 80% ethanol and vortex it:

###### Fresh 80% ethanol

reactions: 

|  |  |
| --- | --- |
| ethanol | mL |
| water | mL |

total:  mL

**Note:** Certified molecular biology-grade water is not required for the 80% ethanol. Low-retention
pipet tips are not required for the next steps, until resuspending the pellet.

**Note:** Perform the remaining steps quickly, until the pellet is resuspended, to prevent
the beads from overdrying. If you have a large number of samples, perform the next
steps on only half of them before coming back to this point for the other half.

When the beads have fully separated, carefully remove and discard the **~28 µL** supernatants without disturbing the bead pellets (pipet slowly from the side opposite
the magnet).

Add **200 µL fresh 80% ethanol** to each sample.

Carefully remove and discard the **~200 µL** supernatants without disturbing the bead pellets.

Briefly centrifuge the samples to collect any remaining ethanol droplets.

Return the samples to the magnet and carefully remove and discard all remaining supernatant
with a smaller pipet.

Away from the magnet, resuspend each bead pellet in **10 µL DNA storage buffer** by **pipetting to mix** until the bead clumps are broken up and evenly mixed.

Briefly centrifuge the samples to remove bubbles, then return them to the magnet to
separate the beads. They should separate completely within a few seconds.

Transfer the **~10 µL** supernatants to new 0.2 mL low-retention tubes. These are your sequencing-ready libraries.
Keep them refrigerated for short-term use, to reduce freezing and thawing, or frozen
for long-term storage.

---

### Next steps

#### Quantification and validation of the libraries

The final yield should be 10 μL of amplified library between 50 and 200 nM, with most
of the fragments between 150 and 400 bp. The size distribution can be verified by
running the library undiluted with a TapeStation High Sensitivity D1000 kit or equivalent.
The electropherogram may show evidence of overamplification: a secondary bump or especially
wide smear of molecules that migrate more slowly than the rest, because they comprise
complementary annealed adapters and noncomplementary, unannealable inserts. These
libraries can still be sequenced, but this artifact makes molarity and size-distribution
estimates inaccurate, and it is ideal to recalibrate the PCR cycles to the maximum
number that does not produce this artifact.

To determine pooling volumes (if using cleanup for individual samples) and optimize loading concentrations, it is helpful to measure the concentration
more precisely by qPCR, which measures sequenceable molecules rather than total DNA
content. Roche's KAPA Library Quantification Kits are designed for this purpose.

#### Sequencing

A good minimum target for the human genome is at least 40 million read pairs per library,
e.g. up to 96 libraries on an Illumina NovaSeq S2 flow cell. Long reads (> 50 nt)
are not useful as this method counts fragments, not bases, but paired-end reads are
useful to count both ends of each fragment (use 2x50 on NovaSeq and NextSeq 1000/200,
2x38 on NextSeq 500). Be sure to include the correct index read lengths for your indexing scheme (8 nt i7 + 8 nt i5 for combinatorial dual indexing, 10 nt i7 + no i5 for i7-only
unique single indexing based on IDT/Illumina UDIs). No custom sequencing primers are
required as the libraries have standard Nextera adapters. ΦX174 spike-in is not required,
but a 1% spike-in is recommended in case the instrument fails and troubleshooting
is required.

From the MspJI restriction motif, more than 50% of bases at position 17 in both reads
will be G and more than 50% at position 14 will be C or T. This does not interfere
with sequencing performance. The organism's methylation motif will also be reflected
here, e.g. a human genome with CpG methylation will also be more than 50% C at position
16. GC content throughout the reads may also be higher than across the entire genome
if DNA methylation is enriched in GC-rich regions, as in the human genome.

#### Data processing

The sequence reads can be aligned to a reference genome with any standard aligner.
The restriction digestion fragments DNA at a small number of specific positions, so
duplicate fragments will frequently occur by chance; **do not remove duplicates**.

Suggested pipeline (implemented in FMLtools):

1. Demultiplex pooled libraries with Illumina software.
2. Trim Nextera adapter sequences with CutAdapt.
3. Align reads to the reference genome with bwa-mem2.
4. Sort and index alignments with Samtools.
5. Count hits per motif site with `count_fml_hits.py`.
6. Aggregate hits per genome region with `region_counts.py`.
7. Analyze read counts with DESeq2.

Required one-time preparation:

1. Index the reference genome sequence with the aligner.
2. Index motif sites with `get_sequence_positions.py`.
3. Obtain a list of genome regions of interest (e.g. promoters) in BED format.

---

### Oligonucleotide designs (Illumina-specific)

Oligonucleotide sequences © 2021 Illumina, Inc. All rights reserved. Derivative works
created by Illumina customers are authorized for use with Illumina instruments and
products only. All other uses are strictly prohibited.

All sequences are in the order 5′→3′ and all nucleotides are DNA. All cytosines are
unmethylated.

#### Abbreviations

- `N`: equimolar mix of A, C, G, T
- `U`: deoxyuridine (DNA not RNA)
- `*`: phosphorothioate bond
- `[index]`: distinct index sequence for each version of the primer

#### Library synthesis adapters

Use HPLC purification. This reduces the complexity of the random bases, but that is
less important than the purity of full-length molecules. These oligos form secondary
structures in moderate salt conditions but that should not affect spectrophotometry
in storage buffer. Although they are used for ligation, the adapters **must not have a 5′ phosphate**; synthesis without it is standard so simply do not request the optional 5′ phosphate.

- FML-seq P5 adapter: `NNNNCUGUCUCUUAUACACAUCUTCGTCGGCAGCGTCAGATGTGTATAAGAGACAG`
- FML-seq P7 adapter: `NNNNCUGUCUCUUAUACACAUCUGTCTCGTGGGCTCGGAGATGTGTATAAGAGACAG`

#### Indexing PCR primers

Use IDT's Ultramer synthesis or HPLC purification. Choose any indexing scheme and
order Nextera-compatible DNA primers using the provided designs, keeping most of the
sequence constant and varying only the index. Combinatorial dual indexing is inexpensive
but requires careful arrangement of the index pairs. Alternatively, use a P5 primer with no index and only the i7 indexes from the IDT/Illumina
unique dual indexing scheme for Nextera, which is more expensive but less difficult
to prepare and saves sequencing cycles that can be used to increase the read lengths
for the insert instead. Unique dual indexing is not likely to be helpful for FML-seq.
**TruSeq primers are not compatible** and index sequences validated with TruSeq adapters are not recommended for these
Nextera adapters.

- Nextera P5 PCR primer: `AATGATACGGCGACCACCGAGATCTACAC[index]TCGTCGGCAGCGT*C`
- Nextera P7 PCR primer: `CAAGCAGAAGACGGCATACGAGAT[index]GTCTCGTGGGCTCG*G`

---

### Quick reference protocol

reactions: 

#### Digestion

###### Digestion Mix: 4 µL + 1 µL gDNA, then start Program 1

|  |  |
| --- | --- |
| water | µL |
| rCutSmart Buffer, 10X | µL |
| premade Oligo Mix | µL |
| MspJI, 5 U/µL | µL |

total:  µL

#### Synthesis

###### Synthesis Mix: add 4 µL + 1 µL primers, then start Program 2

|  |  |
| --- | --- |
| premade Synthesis Buffer | µL |
| premade Synthesis Enzymes | µL |

total:  µL

#### Cleanup

Mix with **18 µL SPRI bead mix**.

Incubate 1 min, then centrifuge.

Separate on magnet and discard supernatant.

Wash with **200 µL fresh 80% ethanol**, then discard supernatant.

###### Fresh 80% ethanol

|  |  |
| --- | --- |
| ethanol | mL |
| water | mL |

total:  mL

Centrifuge, then discard supernatant again.

Resuspend in **10 µL DNA Storage Solution**, then briefly centrifuge.

Separate on magnet and transfer libraries to new tubes.
