## Supplemental Code for "Cost-effective DNA methylation profiling by FML-seq": element_density_fml.R

```
#! /usr/bin/Rscript

library(ggplot2)
library(scales)
library(parallel)
options(mc.cores = detectCores())

# nice lookup table of names
annotations <- c(
	refseq_curated_exons = "Exons",
	refseq_curated_introns = "Introns",
	ENCODE_cCREs = "Promoters",
	refseq_enhancers = "Enhancers"
)


motif.counts <- setNames(mclapply(names(annotations), function(name) {
	result.raw <- scan(paste0(name, ".CGNR.tsv"), list(character(), integer()), quiet = T)
	setNames(result.raw[[2]], result.raw[[1]])
}), names(annotations))

rpmpm <- do.call(rbind, mclapply(Sys.glob("hit_counts_fml/*.tsv"), function(file) {
	library <- sub("\\..*", "", basename(file))
	cell <- sub("_.*", "", basename(file))
	annotation <- sub(".*\\.", "", sub("\\.tsv$", "", file))
	counts <- scan(file, list(NULL, integer()), quiet = T)[[2]]
	data.frame(
		library,
		cell,
		annotation,
		rpmpm = counts / motif.counts[[annotation]] / sum(counts) * 1E6
	)
}))
rpmpm$nice.annotation <- factor(annotations[rpmpm$annotation], levels = annotations)

plot.fml.density <- ggplot(rpmpm, aes(library, rpmpm, fill = cell)) +
	geom_violin() +
	xlab("Technical-duplicate FML-seq libraries (60 ng gDNA input)") +
	scale_y_log10(name = "FML-seq relative density (RPMPM)", label = comma) +
	facet_grid(~ nice.annotation) +
	theme(
		axis.ticks.x = element_blank(),
		axis.text.x = element_blank()
	)

ggsave(plot.fml.density, file = "element_density_fml.pdf", width = 8, height = 3)

save.image("element_density_fml.RData")
```
