## Supplemental Code for "Cost-effective DNA methylation profiling by FML-seq": element_pmethyl_wgbs.R

```
#! /usr/bin/Rscript

library(ggplot2)
library(scales)
library(parallel)
options(mc.cores = detectCores())

# nice lookup table of names
annotations <- c(
	refseq_curated_exons = "Exons",
	refseq_curated_introns = "Introns",
	ENCODE_cCREs = "Promoters",
	refseq_enhancers = "Enhancers"
)

pmethyl <- do.call(rbind, mclapply(Sys.glob("hit_counts_wgbs/*.tsv"), function(file) {
	library <- sub("\\..*", "", basename(file))
	cell <- sub("_.*", "", basename(file))
	annotation <- sub(".*\\.", "", sub("\\.tsv$", "", file))
	counts.raw <- scan(file, list(NULL, integer(), integer()), quiet = T)
	data.frame(
		library,
		cell,
		annotation,
		pmethyl = counts.raw[[2]] / (counts.raw[[2]] + counts.raw[[3]])
	)
}))
pmethyl$nice.annotation <- factor(annotations[pmethyl$annotation], levels = annotations)

plot.wgbs.pmethyl <- ggplot(pmethyl, aes(library, pmethyl, fill = cell)) +
	geom_violin() +
	xlab("Technical-duplicate WGBS libraries") +
	scale_y_continuous(name = "Methylation (base nonconversion)", label = percent) +
	facet_grid(~ nice.annotation) +
	theme(
		axis.ticks.x = element_blank(),
		axis.text.x = element_blank()
	)

ggsave(plot.wgbs.pmethyl, file = "element_pmethyl_wgbs.pdf", width = 8, height = 3)

save.image("element_pmethyl_wgbs.RData")
```
