## Supplemental Code for "Cost-effective DNA methylation profiling by FML-seq": test_length.R

```
#! /usr/bin/Rscript

library(ggplot2)
library(scales)

totals.raw <- read.delim("bwa_stats.tsv")
totals <- setNames(totals.raw$total.reads, totals.raw$library)

counts <- read.delim("test_length.tsv")
counts$cell <- sapply(counts$library, function(x) strsplit(x, "_")[[1]][1])
counts$prop <- counts$confidently.aligned / totals[counts$library]

pdf("test_length.pdf", width = 4, height = 3)
ggplot(counts, aes(length, prop, color = cell, group = library)) +
	geom_line() +
	scale_x_continuous(name = "sequence read length, each end (nt)") +
	scale_y_continuous(name = "confidently aligned reads", label = percent, limits = c(0, 1)) +
	geom_vline(xintercept = c(38, 51, 76, 101, 151), linetype = 2, color = "darkgray") +
	theme(legend.position = c(1, 0), legend.justification = c(1, 0))
dev.off()
```
