## Supplemental Code for "Cost-effective DNA methylation profiling by FML-seq": test_length.sh

```
#! /bin/bash

set -euo pipefail

echo -e "library\tlength\tconfidently aligned" > test_length.tsv
parallel -k '
	fastq1=$(readlink -f {2})
	fastq2=$(sed "s/_R1\.fastq\.gz$/_R2.fastq.gz/" <<< $fastq1)
	library_name=$(basename {2} _R1.fastq.gz)
	echo -en "$library_name\t{1}\t"
	
	$HOME/Downloads/novocraft3/novoalign -n {1} -F ILM1.8 -a CTGTCTCTTATACACATCTCCGAGCCCACGAGAC CTGTCTCTTATACACATCTGACGCTGCCGACGA -o SAM -d $HOME/genome/hs1/hs1.ndx -f <(zstd -dc $fastq1) <(zstd -dc $fastq2) 2> /dev/null |
		samtools view -cq 10
	
	echo -e "aligned $library_name with length {1}\n" >&2
' ::: {25..154} ::: fastq/*60ng*_R1.fastq.gz >> test_length.tsv
```
