## Supplemental Code for "Cost-effective DNA methylation profiling by FML-seq": deseq_2conditions.R

```
#! /usr/bin/Rscript

library(DESeq2)

CELLS <- c("K562", "HeLa-S3") # use only these cell types for a two-class comparison

load("counts_60ng.RData")

which.include <- which(design.fml$cell %in% CELLS)
dds.2conditions <- DESeq(DESeqDataSetFromMatrix(counts.fml[,which.include], colData = design.fml[which.include,], design = ~ cell), fitType = "local")
results.2conditions <- results(dds.2conditions, parallel = T)

save(dds.2conditions, results.2conditions, file = "deseq_2conditions.RData")
```
