## Supplemental Code for "Cost-effective DNA methylation profiling by FML-seq": dmr_2conditions_bs.R

```
#! /usr/bin/Rscript

library(parallel)
library(methylKit)

TMP.DIR <- "/tmp/"
CELLS <- c("K562", "HeLa-S3") # use only these cell types for a two-class comparison

load("wgbs.RData")

# simulate input files with each promoter as a chromosome
for (library in subset(design.wgbs, cell %in% CELLS)$sample) {
	write.table(data.frame(
		chrBase = "NA",
		chr = dimnames(counts.wgbs)[[1]],
		start = 1,
		strand = "+",
		coverage = rowSums(counts.wgbs[,,library]),
		freqC = 100 * pmethyl.wgbs[,library],
		freqT = 100 * (1 - pmethyl.wgbs[,library])
	), file = paste0(TMP.DIR, library, ".tsv"), sep = "\t", quote = F, row.names = F)
}

meth.wgbs <- unite(methRead(
	as.list(paste0(TMP.DIR, subset(design.wgbs, cell %in% CELLS)$sample, ".tsv")),
	sample.id = as.list(subset(design.wgbs, cell %in% CELLS)$sample),
	assembly = "dummy",
	treatment = factor(subset(design.wgbs, cell %in% CELLS)$cell)
))

diff.wgbs <- calculateDiffMeth(meth.wgbs, mc.cores = detectCores())

# initialize result with NA placeholders
dmr.2conds.wgbs <- data.frame(
	row.names = dimnames(counts.wgbs)[[1]],
	pval = rep(NA, dim(counts.wgbs)[1]),
	padj = NA
)
dmr.2conds.wgbs[diff.wgbs$chr,] <- cbind(diff.wgbs$pvalue, p.adjust(diff.wgbs$pvalue, "BH"))


load("rrbs.RData")

# simulate input files with each promoter as a chromosome
for (library in subset(design.rrbs, cell %in% CELLS)$sample) {
	write.table(data.frame(
		chrBase = "NA",
		chr = dimnames(counts.rrbs)[[1]],
		start = 1,
		strand = "+",
		coverage = rowSums(counts.rrbs[,,library]),
		freqC = 100 * pmethyl.rrbs[,library],
		freqT = 100 * (1 - pmethyl.rrbs[,library])
	), file = paste0(TMP.DIR, library, ".tsv"), sep = "\t", quote = F, row.names = F)
}

meth.rrbs <- unite(methRead(
	as.list(paste0(TMP.DIR, subset(design.rrbs, cell %in% CELLS)$sample, ".tsv")),
	sample.id = as.list(subset(design.rrbs, cell %in% CELLS)$sample),
	assembly = "dummy",
	treatment = factor(subset(design.rrbs, cell %in% CELLS)$cell)
))

diff.rrbs <- calculateDiffMeth(meth.rrbs, mc.cores = detectCores())

# initialize result with NA placeholders
dmr.2conds.rrbs <- data.frame(
	row.names = dimnames(counts.rrbs)[[1]],
	pval = rep(NA, dim(counts.rrbs)[1]),
	padj = NA
)
dmr.2conds.rrbs[diff.rrbs$chr,] <- cbind(diff.rrbs$pvalue, p.adjust(diff.rrbs$pvalue, "BH"))


save(dmr.2conds.wgbs, dmr.2conds.rrbs, file = "dmr_2conditions_bs.RData")
```
