## Supplemental Code for "Cost-effective DNA methylation profiling by FML-seq": fml_by_position.R

```
#! /usr/bin/Rscript

library(abind)
library(ggplot2)
library(scales)
library(DESeq2)
library(parallel)
options(mc.cores = detectCores())

CELLS <- c("K562", "HeLa-S3") # use only these cell types for a two-class comparison

positions.raw <- read.table("CG_hs1_chr22_cpos.bed")
positions.integer <- positions.raw[,3] # right-end positions of BED features
positions <- paste0(positions.raw[,1], ":", positions.raw[,3])
rm(positions.raw, positions.integer) # free up some memory

has.motif <- as.logical(scan("CGNR_present.tsv", integer(), quiet = T))

files.fml <- Sys.glob("*60ng*.chr22_pos_hits.tsv")
counts.fml <- do.call(cbind, mclapply(files.fml, scan, integer(), quiet = T))
colnames(counts.fml) <- sub("\\..*", "", files.fml)
rpm.fml <- counts.fml / colSums(counts.fml)
rpm.fml[! has.motif,] <- NA

cell.fml <- sub("_.*", "", colnames(counts.fml))
counts.pooled.fml <- do.call(cbind, mclapply(unique(cell.fml), function(this.cell) rowSums(counts.fml[,cell.fml == this.cell])))
colnames(counts.pooled.fml) <- unique(cell.fml)
rpm.pooled.fml <- counts.pooled.fml / colSums(counts.pooled.fml) * 1E6
rpm.pooled.fml[! has.motif,] <- NA

files.wgbs <- Sys.glob("*WGBS*.chr22_pos_hits.tsv")
counts.wgbs <- do.call(abind, c(mclapply(files.wgbs, read.table), along = 3))
dimnames(counts.wgbs)[[2]] <- c("methylated", "unmethylated")
dimnames(counts.wgbs)[[3]] <- sub("\\..*", "", files.wgbs)
cell.wgbs <- sub("_.*", "", files.wgbs)
counts.pooled.wgbs <- abind(mclapply(unique(cell.wgbs), function(this.cell) apply(counts.wgbs[,, cell.wgbs == this.cell], c(1, 2), sum)), along = 3, new.names = unique(cell.wgbs))
pmethyl.pooled.wgbs <- counts.pooled.wgbs[,1,] / apply(counts.pooled.wgbs, c(1, 3), sum)

beta.by.probe <- read.table("beta_by_probe.tsv", check.names = F)
probe.positions <- read.table("infinium-methylationepic_hs1.CG_only.bed")
probe.positions <- subset(probe.positions, V1 == "chr22")
probe.lookup <- setNames(paste0("chr22:", probe.positions$V3 - (probe.positions$V6 == "+")), probe.positions$V4)
beta.by.probe <- beta.by.probe[rownames(beta.by.probe) %in% names(probe.lookup),]
beta.by.pos <- matrix(as.numeric(NA), nrow = length(positions), ncol = ncol(beta.by.probe), dimnames = list(positions, colnames(beta.by.probe)))
for (probe in rownames(beta.by.probe)) for (col in colnames(beta.by.probe)) beta.by.pos[probe.lookup[probe],col] <- beta.by.probe[probe,col] # inefficient, sorry

which.include <- which(cell.fml %in% CELLS)
dds.2conditions <- DESeq(DESeqDataSetFromMatrix(counts.fml[,which.include], colData = data.frame(cell = cell.fml[which.include]), design = ~ cell), fitType = "local")
results.2conditions <- results(dds.2conditions, parallel = T)
foldchange.fml <- 2^(-results.2conditions$log2FoldChange) # reverse sign because test was backwards
foldchange.wgbs <- pmethyl.pooled.wgbs[,CELLS[2]] / pmethyl.pooled.wgbs[,CELLS[1]]
foldchange.epic <- beta.by.pos[,CELLS[2]] / beta.by.pos[,CELLS[1]]


random.subsample <- sample(which(
	rowSums(is.na(rpm.pooled.fml)) == 0 &
	rowSums(is.na(pmethyl.pooled.wgbs)) == 0 &
	rowSums(is.na(beta.by.pos)) == 0
), 200)


save.image("fml_by_position.RData")


plot.fml.density <- qplot(as.vector(counts.fml / has.motif), xlim = c(NA, 100)) +
	xlab("Reads assigned to cytosine position") +
	scale_y_continuous(name = "Cytosine positions", label = comma)
ggsave(plot.fml.density, file = "position_density_fml.pdf", width = 4, height = 2)

pdf("position_scatterplot_2conditions_score.pdf", width = 2.75, height = 2)
qplot(pmethyl.pooled.wgbs[random.subsample,CELLS[1]], rpm.pooled.fml[random.subsample,CELLS[1]], main = "Methylation score") +
	scale_x_continuous(label = percent, name = "WGBS (% unconverted)") +
	scale_y_continuous(label = comma, name = "FML-seq (reads per million)") +
	annotate("text", x = 0, y = Inf, label = bquote(rho == .(round(cor(pmethyl.pooled.wgbs[,CELLS[1]], rpm.pooled.fml[,CELLS[1]], method = "spearman", use = "complete.obs"), 3))), hjust = 0, vjust = 1)
qplot(pmethyl.pooled.wgbs[random.subsample,CELLS[1]], beta.by.pos[random.subsample,CELLS[1]], main = "Methylation score") +
	scale_x_log10(label = percent, name = "WGBS (% unconverted)") +
	scale_y_log10(label = comma, name = expression(paste("EPIC (norm. ", beta, ")"))) +
	geom_smooth(method = "lm") +
	annotate("text", x = Inf, y = 0, label = bquote(rho == .(round(cor(pmethyl.pooled.wgbs[,CELLS[1]], beta.by.pos[,CELLS[1]], method = "spearman", use = "complete.obs"), 3))), hjust = 1, vjust = 0)
dev.off()

pdf("position_hexplot_2conditions_score.pdf", width = 3.25, height = 2)
qplot(pmethyl.pooled.wgbs[,CELLS[1]], rpm.pooled.fml[,CELLS[1]], main = "Methylation score", geom = "hex") +
	scale_x_continuous(label = percent, name = "WGBS (% unconverted)") +
	scale_y_continuous(label = comma, name = "FML-seq (reads per million)") +
	scale_fill_gradient(name = "C positions", trans = "log", breaks = 10^(0:7), label = comma) +
	annotate("text", x = 0, y = Inf, label = bquote(rho == .(round(cor(pmethyl.pooled.wgbs[,CELLS[1]], rpm.pooled.fml[,CELLS[1]], method = "spearman", use = "complete.obs"), 3))), hjust = 0, vjust = 1)
qplot(pmethyl.pooled.wgbs[,CELLS[1]], beta.by.pos[,CELLS[1]], main = "Methylation score", geom = "hex") +
	scale_x_log10(label = percent, name = "WGBS (% unconverted)") +
	scale_y_log10(label = comma, name = expression(paste("EPIC (norm. ", beta, ")"))) +
	scale_fill_gradient(name = "C positions", trans = "log", breaks = 10^(0:7), label = comma) +
	annotate("text", x = 0, y = Inf, label = bquote(rho == .(round(cor(pmethyl.pooled.wgbs[,CELLS[1]], beta.by.pos[,CELLS[1]], method = "spearman", use = "complete.obs"), 3))), hjust = 0, vjust = 1)
dev.off()


pdf("position_scatterplot_2conditions_foldchange.pdf", width = 2.75, height = 2)
qplot(foldchange.wgbs[random.subsample], foldchange.fml[random.subsample], main = "Differential methylation") +
	scale_x_log10(label = comma, name = "WGBS (% unconverted ratio)") +
	scale_y_log10(label = comma, name = "FML-seq (DESeq2)") +
	geom_abline(slope = 1, color = "red") +
	annotate("text", x = Inf, y = 0, label = bquote(rho == .(round(cor(foldchange.wgbs, foldchange.fml, method = "spearman", use = "complete.obs"), 3))), hjust = 1, vjust = 0)
qplot(foldchange.wgbs[random.subsample], foldchange.epic[random.subsample], main = "Differential methylation") +
	scale_x_log10(label = comma, name = "WGBS (% unconverted ratio)") +
	scale_y_log10(label = comma, name = expression(paste("EPIC (norm. ", beta, " ratio)"))) +
	geom_abline(slope = 1, color = "red") +
	annotate("text", x = Inf, y = 0, label = bquote(rho == .(round(cor(foldchange.wgbs, foldchange.epic, method = "spearman", use = "complete.obs"), 3))), hjust = 1, vjust = 0)
dev.off()

pdf("position_hexplot_2conditions_foldchange.pdf", width = 3.25, height = 2)
qplot(foldchange.wgbs, foldchange.fml, main = "Differential methylation", geom = "hex") +
	scale_x_log10(label = comma, name = "WGBS (% unconverted ratio)") +
	scale_y_log10(label = comma, name = "FML-seq (DESeq2)") +
	scale_fill_gradient(name = "C positions", trans = "log", breaks = 10^(0:7), label = comma) +
	geom_abline(slope = 1, color = "red") +
	annotate("text", x = Inf, y = 0, label = bquote(rho == .(round(cor(foldchange.wgbs, foldchange.fml, method = "spearman", use = "complete.obs"), 3))), hjust = 1, vjust = 0)
qplot(foldchange.wgbs, foldchange.epic, main = "Differential methylation", geom = "hex") +
	scale_x_log10(label = comma, name = "WGBS (% unconverted ratio)") +
	scale_y_log10(label = comma, name = expression(paste("EPIC (norm. ", beta, " ratio)"))) +
	scale_fill_gradient(name = "C positions", trans = "log", breaks = 10^(0:7), label = comma) +
	geom_abline(slope = 1, color = "red") +
	annotate("text", x = 0, y = Inf, label = bquote(rho == .(round(cor(foldchange.wgbs, foldchange.epic, method = "spearman", use = "complete.obs"), 3))), hjust = 0, vjust = 1)
dev.off()
```
