## Supplemental Code for "Cost-effective DNA methylation profiling by FML-seq": heatmap_top500.R

```
#! /usr/bin/Rscript

library(ggplot2)
library(scales)

matnorm <- function(mat) (mat - min(mat, na.rm = T)) / diff(range(mat, na.rm =T))

load("epic.RData")
load("wgbs.RData")
load("rrbs.RData")
load("medip.RData")
load("counts_60ng.RData")


which.all.methods <- which(
	rowSums(is.na(beta.epic)) == 0 &
	rowSums(is.na(pmethyl.wgbs)) == 0 &
	rowSums(is.na(pmethyl.rrbs)) == 0 &
	rowSums(counts.medip) > 0 &
	rowSums(counts.fml) > 0
)


# heatmap of top 500 most variable promoters measured on all platforms
chisq.results.wgbs <- t(apply(counts.pooled.wgbs[which.all.methods,,], 1, function(mat) unlist(chisq.test(mat)[c("statistic", "p.value")]))) # this is bad please improve
top500 <- order(chisq.results.wgbs[,"statistic.X-squared"], na.last = T, decreasing = T)[1:500]

matnorm.combined.top500 <- do.call(cbind, lapply(list(beta.epic, pmethyl.wgbs, pmethyl.rrbs, vsd.medip, log10rpmpm.fml), function(mat) matnorm(mat[which.all.methods,][top500,])))

pdf("heatmap_top500.pdf", width = 7.5, height = 7.5) # it's fussy about the dimensions
	heatmap(matnorm.combined.top500,
		scale = "none",
		distfun = function(mat) as.dist(1 - cor(t(mat))), # "Pearson distance"
		hclustfun = function(mat) hclust(mat, method = "average"), # UPGMA
		col = colorRampPalette(c("white", "darkblue"))(64),
		labRow = NA,
		labCol = colnames(matnorm.combined.top500), # trim the replicate names 
		margins = c(10, 1)
	)
dev.off()
```
