## Supplemental Code for "Cost-effective DNA methylation profiling by FML-seq": load_60ng.R

```
#! /usr/bin/Rscript

library(parallel)
options(mc.cores = detectCores())

promoter.contents <- read.delim("~/genome/hs1/ENCODE_hs1_cCREs_promoter-like.150-350bp.contents.tsv")

files <- Sys.glob("promoter_hit/*60ng*.tsv")
design.fml <- data.frame(
	sample = sub("60ng", "FML-seq", sub("\\..*", "", basename(files)))
)
design.fml$cell <- sapply(design.fml$sample, function(name) strsplit(name, "_")[[1]][1])
design.fml$method <- "FML-seq"

counts.fml <- do.call(cbind, mclapply(files, function(file) as.matrix(read.delim(file, header = F, row.names = 1))))
colnames(counts.fml) <- design.fml$sample

rpm.fml <- sweep(counts.fml, 2, colSums(counts.fml), "/") * 1E6
rpmpm.fml <- sweep(rpm.fml, 1, promoter.contents$CGNR, "/")
log10rpmpm.fml <- log10(rpmpm.fml + 1)

counts.pooled.fml <- do.call(cbind, mclapply(unique(design.fml$cell), function(this.cell.type) rowSums(counts.fml[,design.fml$cell == this.cell.type])))
colnames(counts.pooled.fml) <- unique(design.fml$cell)
rpm.pooled.fml <- sweep(counts.pooled.fml, 2, colSums(counts.pooled.fml), "/") * 1E6
rpmpm.pooled.fml <- sweep(rpm.pooled.fml, 1, promoter.contents$CGNR, "/")
log10rpmpm.pooled.fml <- log10(rpmpm.pooled.fml + 1)


save(design.fml, counts.fml,rpmpm.fml, log10rpmpm.fml, counts.pooled.fml, rpmpm.pooled.fml, log10rpmpm.pooled.fml, file = "counts_60ng.RData")
```
