## Supplemental Code for "Cost-effective DNA methylation profiling by FML-seq": load_epic.R

```
#! /usr/bin/Rscript

# extract beta values from idat and export into (unsorted, hg19) BED
# beta is encoded as the "score" BED field, multiplied by 1000 because it's supposed to range from 0 to 1000

library(parallel)
library(ChAMP)

options(mc.cores = detectCores(), scipen = 999) # disable scientific notation in output

combine.probes <- function(mat.by.probe, lookup.list, combine.fun = colMeans, ...) do.call(rbind, mclapply(lookup.list, function(probes) {
	sublist <- intersect(probes, rownames(mat.by.probe))
	if (length(sublist) == 0) {
		rep(NA, ncol(mat.by.probe))
	} else {
		submat = mat.by.probe[sublist,]
		if (length(sublist) == 1) {
			submat
		} else {
			if (ncol(mat.by.probe) == 1) submat <- as.matrix(mat.by.probe[sublist,]) # keep matrix as matrix
			combine.fun(submat, ...)
		}
	}
}))


# read promoters
promoter.probes.raw <- scan("~/genome/hs1/ENCODE_hs1_cCREs_promoter-like.150-350bp.epic_CG.contents.tsv", list(character(), character()), fill = T)
promoter.probes <- setNames(strsplit(promoter.probes.raw[[2]], ","), promoter.probes.raw[[1]])

# do ChAMP
myLoad <- champ.load("epic/mock_dir", arraytype = "EPIC")
betanorm <- champ.norm(myLoad$beta, cores = detectCores())
write.table(betanorm, "epic/beta_by_probe.tsv", sep = "\t", quote = F, col.names = NA)

# get combined data by promoter (takes a long time and a lot of memory!)
beta.epic <- combine.probes(betanorm, promoter.probes)
colnames(beta.epic) <- paste0(colnames(beta.epic), "_EPIC")

save(beta.epic, file = "epic.RData")
```
