## Supplemental Code for "Cost-effective DNA methylation profiling by FML-seq": load_medip.R

```
#! /usr/bin/Rscript

library(DESeq2)

library(parallel)
options(mc.cores = detectCores())

files <- Sys.glob("medip/cram/*.ENCODE_cCREs.tsv")
design.medip <- data.frame(
	sample = sub("\\..*", "_MeDIP", basename(files))
)
design.medip$cell <- sapply(design.medip$sample, function(name) strsplit(name, "_")[[1]][1])
design.medip$method <- "MeDIP"

counts.medip <- do.call(cbind, mclapply(files, function(file) as.matrix(read.delim(file, header = F, row.names = 1))))
colnames(counts.medip) <- paste(design.medip$cell, design.medip$method, sep = "_")

dds.medip <- DESeq(DESeqDataSetFromMatrix(counts.medip, colData = data.frame(cell = design.medip$cell), design = ~ 1), fitType = "local", parallel = T) # design set to intercept because there are no replicates
vsd.medip <- assay(varianceStabilizingTransformation(dds.medip))

save(design.medip, counts.medip, dds.medip, vsd.medip, file = "medip.RData")
```
