## Supplemental Code for "Cost-effective DNA methylation profiling by FML-seq": load_rrbs.R

```
#! /usr/bin/Rscript

library(abind)
library(parallel)
options(mc.cores = detectCores())

files <- Sys.glob("rrbs/cram/*_CpG.ENCODE_cCREs.tsv")
design.rrbs <- data.frame(
	sample = sub("_CpG.*", "", basename(files))
)
design.rrbs$cell <- sapply(design.rrbs$sample, function(name) strsplit(name, "_")[[1]][1])
design.rrbs$method <- "RRBS"

counts.rrbs <- abind(mclapply(files, function(file) as.matrix(read.delim(file, header = F, row.names = 1, col.names = c(NA, "methylated", "unmethylated")))), along = 3, new.names = design.rrbs$sample)

totals.rrbs <- apply(counts.rrbs, c(1, 3), sum)
pmethyl.rrbs <- counts.rrbs[,1,] / totals.rrbs

save(design.rrbs, counts.rrbs, pmethyl.rrbs, file = "rrbs.RData")
```
