## Supplemental Code for "Cost-effective DNA methylation profiling by FML-seq": load_wgbs.R

```
#! /usr/bin/Rscript

library(abind)
library(parallel)
options(mc.cores = detectCores())

files <- Sys.glob("wgbs/cram/*_CpG.ENCODE_cCREs.tsv")
design.wgbs <- data.frame(
	sample = sub("_CpG.*", "", basename(files))
)
design.wgbs$cell <- sapply(design.wgbs$sample, function(name) strsplit(name, "_")[[1]][1])
design.wgbs$method <- "WGBS"

counts.wgbs <- abind(mclapply(files, function(file) as.matrix(read.delim(file, header = F, row.names = 1, col.names = c(NA, "methylated", "unmethylated")))), along = 3, new.names = design.wgbs$sample)

counts.pooled.wgbs <- abind(mclapply(unique(design.wgbs$cell), function(this.cell.type) apply(counts.wgbs[,, design.wgbs$cell == this.cell.type], c(1, 2), sum)), along = 3, new.names = unique(design.wgbs$cell))

totals.wgbs <- apply(counts.wgbs, c(1, 3), sum)
pmethyl.wgbs <- counts.wgbs[,1,] / totals.wgbs
pmethyl.pooled.wgbs <- counts.pooled.wgbs[,1,] / apply(counts.pooled.wgbs, c(1, 3), sum)

save(design.wgbs, counts.wgbs, counts.pooled.wgbs, pmethyl.wgbs, pmethyl.pooled.wgbs, file = "wgbs.RData")
```
