## Supplemental Code for "Cost-effective DNA methylation profiling by FML-seq": region_dackel.py

```
#! /usr/bin/env -S pypy3 -OO

import click
import genomerator as gen

def zero_data (feature):
	feature.data['counts'] = [0, 0]
	return feature

def update_region (region, position):
	for i in (0, 1): region.data['counts'][i] += int(position.data[i + 1]) # assumes MethylDackel bedGraph format

@click.command()
@click.option('-z', '--zeroes', is_flag = True, help = 'output regions with zero count')
@click.option('-q', '--quiet', is_flag = True, help = 'don\'t show progress bar')
@click.argument('reference_file', type = click.File('r'))
@click.argument('region_bed', type = click.File('r'))
@click.argument('count_bedgraph', required = False, type = click.File('r'), default = click.open_file('-', 'r'))
@click.argument('out_tsv', required = False, type = click.File('w'), default = click.open_file('-', 'w'))
def region_dackel (region_bed, count_bedgraph, out_tsv, reference_file, zeroes, quiet):
	'''
	Given a BED file of genome regions, a bedGraph file in MethylDackel format, and a list of reference lengths, count the number of total unmethylated vs. methylated cytosine hits in each genome region.
	Spans in the bedGraph are assumed (not checked!) to be only 1 base each, so they must fall entirely inside the genome region.
	Result is written as three-column TSV with region name, methylated count, unmethylated count.
	'''
	
	references = gen.read_reference_lengths(reference_file)
	reference_lengths = list(references.values())
	region_stream = gen.BedStream(region_bed, assert_sorted = True, references = references.keys())
	count_stream = gen.BedgraphStream(count_bedgraph, assert_sorted = True, parse = False, references = references.keys())
	
	counter = gen.OperationGenerator(
		a = map(zero_data, region_stream),
		b = count_stream,
		match = lambda a,b: b in a,
		a_is_passed = lambda a,b: a.left_of(b),
		b_is_passed = lambda a,b: a.right_of(b),
		operate = update_region
	)
	
	if not quiet: bar = gen.ProgressBar(reference_lengths)
	for region in counter:
		if zeroes or sum(region.data['counts']) > 0: out_tsv.write('\t'.join([region.data['name']] + list(map(str, region.data['counts']))) + '\n')
		if not quiet: bar.update(region)
	if not quiet: bar.finish()

if __name__ == '__main__':
	region_dackel()
```
