## Supplemental Code for "Cost-effective DNA methylation profiling by FML-seq": region_medip.py

```
#! /usr/bin/env -S python3 -OO

import click
import pysam
import genomerator as gen

def zero_data (feature):
	feature.data['count'] = 0
	return feature

def reduce_width (feature, width): # doesn't preserve sortedness so don't map!
	if len(feature) > width:
		feature.left_pos = round((feature.left_pos + feature.right_pos) / 2 - width / 2)
		feature.right_pos = feature.left_pos + width - 1
	return feature

@click.command()
@click.option('-w', '--width', type = int, default = 1, help = 'maximum width to reduce sequenced fragments, at center')
@click.option('-z', '--zeroes', is_flag = True, help = 'output regions with zero count')
@click.option('-q', '--quiet', is_flag = True, help = 'don\'t show progress bar')
@click.argument('reference_file', type = click.File('r'))
@click.argument('region_bed', type = click.File('r'))
@click.argument('medip_sam', required = False, type = str, default = '-')
@click.argument('out_tsv', required = False, type = click.File('w'), default = click.open_file('-', 'w'))
def region_medip (region_bed, medip_sam, out_tsv, reference_file, width, zeroes, quiet):
	'''
	Given a BED file of genome regions, a SAM file of paired-end MeDIP-seq reads, and a list of reference lengths, count the number of reads that hit each region.
	Each MeDIP insert is reduced to the specified width (if not already shorter) at its center,
	then counted as a hit if it intersects a region.
	Result is written as two-column TSV with region name and hit count.
	'''
	
	references = gen.read_reference_lengths(reference_file)
	reference_lengths = list(references.values())
	region_stream = gen.BedStream(region_bed, assert_sorted = True, references = references.keys())
	medip_stream = gen.SamFragmentStream(pysam.Samfile(medip_sam), assert_sorted = True, references = references.keys())
	
	counter = gen.OperationGenerator(
		a = map(zero_data, region_stream),
		b = medip_stream,
		match = lambda a,b: reduce_width(b, width).intersects(a),
		a_is_passed = lambda a,b: a.left_of(b),
		b_is_passed = lambda a,b: a.right_of(b),
		operate = lambda region,feature: region.data.__setitem__('count', region.data['count'] + 1) # increment
	)
	
	if not quiet: bar = gen.ProgressBar(reference_lengths)
	for region in counter:
		if zeroes or region.data['count'] > 0: out_tsv.write('%s\t%i\n' % (region.data['name'], region.data['count']))
		if not quiet: bar.update(region)
	if not quiet: bar.finish()

if __name__ == '__main__':
	region_medip()
```
