## Supplemental Code for "Cost-effective DNA methylation profiling by FML-seq": scatterplots_2conditions.R

```
#! /usr/bin/Rscript

library(ggplot2)
library(scales)

CELLS <- c("K562", "HeLa-S3") # use only these cell types for a two-class comparison

load("counts_60ng.RData")
load("deseq_2conditions.RData")
load("wgbs.RData")
load("epic.RData")

random.subsample <- sample(which(
	rowSums(is.na(rpmpm.pooled.fml)) == 0 &
	rowSums(is.na(pmethyl.pooled.wgbs)) == 0 &
	rowSums(is.na(beta.epic)) == 0
), 200)

colnames(beta.epic) <- sub("_EPIC", "", colnames(beta.epic))

# methylation score
pdf("scatterplots_2conditions_score.pdf", width = 2.75, height = 2)
qplot(pmethyl.pooled.wgbs[random.subsample,CELLS[1]], rpmpm.pooled.fml[random.subsample,CELLS[1]], main = "Methylation score") +
	scale_x_log10(label = percent, name = "WGBS (% unconverted)") +
	scale_y_log10(label = comma, name = "FML-seq (RPMPM)") +
	geom_smooth(method = "lm") +
	annotate("text", x = Inf, y = 0, label = bquote(rho == .(round(cor(pmethyl.pooled.wgbs[,CELLS[1]], rpmpm.pooled.fml[,CELLS[1]], method = "spearman", use = "complete.obs"), 3))), hjust = 1, vjust = 0)
qplot(pmethyl.pooled.wgbs[random.subsample,CELLS[1]], beta.epic[random.subsample,CELLS[1]], main = "Methylation score") +
	scale_x_log10(label = percent, name = "WGBS (% unconverted)") +
	scale_y_log10(label = comma, name = expression(paste("EPIC (norm. ", beta, ")"))) +
	geom_smooth(method = "lm") +
	annotate("text", x = Inf, y = 0, label = bquote(rho == .(round(cor(pmethyl.pooled.wgbs[,CELLS[1]], beta.epic[,CELLS[1]], method = "spearman", use = "complete.obs"), 3))), hjust = 1, vjust = 0)
dev.off()
pdf("hexplots_2conditions_score.pdf", width = 3.25, height = 2)
qplot(pmethyl.pooled.wgbs[,CELLS[1]], rpmpm.pooled.fml[,CELLS[1]], main = "Methylation score", geom = "hex") +
	scale_x_log10(label = percent, name = "WGBS (% unconverted)") +
	scale_y_log10(label = comma, name = "FML-seq (RPMPM)") +
	scale_fill_gradient(name = "promoters", trans = "log", breaks = 10^(0:7), label = comma) +
	annotate("text", x = 0, y = Inf, label = bquote(rho == .(round(cor(pmethyl.pooled.wgbs[,CELLS[1]], rpmpm.pooled.fml[,CELLS[1]], method = "spearman", use = "complete.obs"), 3))), hjust = 0, vjust = 1)
qplot(pmethyl.pooled.wgbs[,CELLS[1]], beta.epic[,CELLS[1]], main = "Methylation score", geom = "hex") +
	scale_x_log10(label = percent, name = "WGBS (% unconverted)") +
	scale_y_log10(label = comma, name = expression(paste("EPIC (norm. ", beta, ")"))) +
	scale_fill_gradient(name = "promoters", trans = "log", breaks = 10^(0:7), label = comma) +
	annotate("text", x = 0, y = Inf, label = bquote(rho == .(round(cor(pmethyl.pooled.wgbs[,CELLS[1]], beta.epic[,CELLS[1]], method = "spearman", use = "complete.obs"), 3))), hjust = 0, vjust = 1)
dev.off()


# fold change
pdf("scatterplots_2conditions_foldchange.pdf", width = 2.75, height = 2)
foldchange.fml <- 2^(-results.2conditions$log2FoldChange) # reverse sign because test was backwards
foldchange.wgbs <- pmethyl.pooled.wgbs[,CELLS[2]] / pmethyl.pooled.wgbs[,CELLS[1]]
foldchange.epic <- beta.epic[,CELLS[2]] / beta.epic[,CELLS[1]]
qplot(foldchange.wgbs[random.subsample], foldchange.fml[random.subsample], main = "Differential methylation") +
	scale_x_log10(label = comma, name = "WGBS (% unconverted ratio)") +
	scale_y_log10(label = comma, name = "FML-seq (DESeq2)") +
	geom_abline(slope = 1, color = "red") +
	annotate("text", x = Inf, y = 0, label = bquote(rho == .(round(cor(foldchange.wgbs, foldchange.fml, method = "spearman", use = "complete.obs"), 3))), hjust = 1, vjust = 0)
qplot(foldchange.wgbs[random.subsample], foldchange.epic[random.subsample], main = "Differential methylation") +
	scale_x_log10(label = comma, name = "WGBS (% unconverted ratio)") +
	scale_y_log10(label = comma, name = expression(paste("EPIC (norm. ", beta, " ratio)"))) +
	geom_abline(slope = 1, color = "red") +
	annotate("text", x = Inf, y = 0, label = bquote(rho == .(round(cor(foldchange.wgbs, foldchange.epic, method = "spearman", use = "complete.obs"), 3))), hjust = 1, vjust = 0)
dev.off()
pdf("hexplots_2conditions_foldchange.pdf", width = 3.25, height = 2)
qplot(foldchange.wgbs, foldchange.fml, main = "Differential methylation", geom = "hex") +
	scale_x_log10(label = comma, name = "WGBS (% unconverted ratio)") +
	scale_y_log10(label = comma, name = "FML-seq (DESeq2)") +
	scale_fill_gradient(name = "promoters", trans = "log", breaks = 10^(0:7), label = comma) +
	geom_abline(slope = 1, color = "red") +
	annotate("text", x = 0, y = Inf, label = bquote(rho == .(round(cor(foldchange.wgbs, foldchange.fml, method = "spearman", use = "complete.obs"), 3))), hjust = 0, vjust = 1)
qplot(foldchange.wgbs, foldchange.epic, main = "Differential methylation", geom = "hex") +
	scale_x_log10(label = comma, name = "WGBS (% unconverted ratio)") +
	scale_y_log10(label = comma, name = expression(paste("EPIC (norm. ", beta, " ratio)"))) +
	scale_fill_gradient(name = "promoters", trans = "log", breaks = 10^(0:7), label = comma) +
	geom_abline(slope = 1, color = "red") +
	annotate("text", x = 0, y = Inf, label = bquote(rho == .(round(cor(foldchange.wgbs, foldchange.epic, method = "spearman", use = "complete.obs"), 3))), hjust = 0, vjust = 1)
dev.off()
```
