## Supplemental Code for "Cost-effective DNA methylation profiling by FML-seq": venn_dmr_2conditions.R

```
#! /usr/bin/Rscript

library(eulerr)

PADJ.THRESHOLD <- 0.01

load("dmr_2conditions_bs.RData")
load("deseq_2conditions.RData")

sig.dmr <- data.frame(
	WGBS = ! (is.na(dmr.2conds.wgbs$padj) | dmr.2conds.wgbs$padj >= PADJ.THRESHOLD),
	RRBS = ! (is.na(dmr.2conds.rrbs$padj) | dmr.2conds.rrbs$padj >= PADJ.THRESHOLD),
	`FML-seq` = ! (is.na(results.2conditions$padj) | results.2conditions$padj >= PADJ.THRESHOLD)
)
print("significant promoters:")
colSums(sig.dmr)

pdf("venn_dmr_2conditions.pdf", width = 4.5, height = 4)
plot(euler(sig.dmr), quantities = T)
dev.off()
```
