## Supplemental Code for "Cost-effective DNA methylation profiling by FML-seq": analyze_subsamples.R

```
#! /usr/bin/Rscript

library(ggplot2)
library(scales)
library(DESeq2)
library(entropy)
library(cluster)
library(parallel)
options(mc.cores = detectCores())

two.cells <- c("HeLa-S3", "IMR-90") # the two that are done at every dilution

# load only the 60 ng samples from first run then combine with second run
load("subsamples_firstrun.RData")
which.60ng <- which(design$amount == "60ng")
counts.firstrun <- counts.list[which.60ng]
design.firstrun <- design[which.60ng,]
log10rpmpm.firstrun <- log10rpmpm.list[which.60ng]
load("subsamples_secondrun.RData")
counts.list <- c(counts.firstrun, counts.list)
design <- rbind(design.firstrun, design)
log10rpmpm.list <- c(log10rpmpm.firstrun, log10rpmpm.list)

# promoters hit at least once
results.by.library <- cbind(design, promoters = unlist(mclapply(counts.list, function(counts) sum(counts > 0))))
plot.promoters.hit <- ggplot(results.by.library, aes(readpairs / 1E6, promoters, color = cell, group = library)) +
	geom_line() +
	facet_grid(amount ~ .) +
	scale_x_continuous(name = "read pairs sequenced per library", label = unit_format(unit = "M", sep = "")) +
	scale_y_continuous(name = "promoters with at least 1 hit", label = comma) +
	theme(legend.position = "n")
ggsave("subsample_promoters_hit.pdf", plot.promoters.hit, width = 3, height = 6)

# entropy of hit counts
results.by.library$entropy <- unlist(mclapply(counts.list, function(counts) entropy(counts, unit = "log2")))
plot.entropy <- ggplot(results.by.library, aes(readpairs / 1E6, entropy, color = cell, group = library)) +
	geom_line() +
	facet_grid(amount ~ .) +
	scale_x_continuous(name = "read pairs sequenced per library", label = unit_format(unit = "M", sep = "")) +
	scale_y_continuous(name = "entropy of promoter hit counts (bits)", limits = c(0, NA)) +
	theme(legend.position = "n")
ggsave("subsample_entropy.pdf", plot.entropy, width = 3, height = 6)

# correlation of technical replicates
results.by.condition <- unique(design[,c("readpairs", "cell", "amount")])
results.by.condition$corr.rep <- unlist(mclapply(1:nrow(results.by.condition), function(i) {
	which.reps <- which(
		design$readpairs == results.by.condition$readpairs[i] &
		design$cell == results.by.condition$cell[i] &
		design$amount == results.by.condition$amount[i]
	)
	stopifnot(length(which.reps) <= 2)
	if (length(which.reps) == 2) cor(log10rpmpm.list[[which.reps[1]]], log10rpmpm.list[[which.reps[2]]], use = "c") else NA
}))
plot.corr.rep <- ggplot(subset(results.by.condition, ! is.na(corr.rep)), aes(readpairs / 1E6, corr.rep, color = cell, group = cell)) +
	geom_line() +
	facet_grid(amount ~ .) +
	scale_x_continuous(name = "read pairs sequenced per library", label = unit_format(unit = "M", sep = "")) +
	scale_y_continuous(name = "correlation of technical replicates", limits = 0:1) +
	theme(legend.position = "n")
ggsave("subsample_corr_rep.pdf", plot.corr.rep, width = 3, height = 6)

# correlation with WGBS
promoter.contents <- read.delim("~/genome/hs1/ENCODE_hs1_cCREs_promoter-like.150-350bp.contents.tsv")
load("wgbs.RData")
pmethyl.rank.pooled.wgbs <- apply(pmethyl.pooled.wgbs, 2, rank, na.last = "keep") # precompute ranks to be reused; R's Spearman correlation uses this configuration of rank()
results.by.condition$corr.wgbs <- unlist(mclapply(1:nrow(results.by.condition), function(i) {
	which.reps <- which(
		design$readpairs == results.by.condition$readpairs[i] &
		design$cell == results.by.condition$cell[i] &
		design$amount == results.by.condition$amount[i]
	)
	stopifnot(length(which.reps) <= 2)
	if (length(which.reps) == 2) {
		counts.pooled <- counts.list[[which.reps[1]]] + counts.list[[which.reps[2]]]
		log10rpmpm.pooled <- log10((counts.pooled / sum(counts.pooled) * 1E6) / promoter.contents$CGNR + 1)
		cor( # Pearson correlation, because we're giving it pre-ranked data (Spearman is Pearson of ranks)
			pmethyl.rank.pooled.wgbs[,results.by.condition$cell[i]],
			rank(log10rpmpm.pooled, na.last = "keep"),
			use = "c"
		)
	} else NA
}))
plot.corr.wgbs <- ggplot(subset(results.by.condition, ! is.na(corr.wgbs)), aes(readpairs / 1E6, corr.wgbs, color = cell, group = cell)) +
	geom_line() +
	facet_grid(amount ~ .) +
	scale_x_continuous(name = "read pairs sequenced per library", label = unit_format(unit = "M", sep = "")) +
	scale_y_continuous(name = "correlation with WGBS % methylation", limits = 0:1) +
	theme(legend.position = "n")
ggsave("subsample_corr_wgbs.pdf", plot.corr.wgbs, width = 3, height = 6)

# DESeq
results.by.dilution <- unique(design[,c("readpairs", "amount")])
results.by.dilution$sig.promoters <- as.integer(unlist(mclapply(1:nrow(results.by.dilution), function(i) { # DESeq seems to randomly fail sometimes and return an error (as character class), so as.integer converts those to NA. we don't need every data point
	which.samples <- which(
		design$readpairs == results.by.dilution$readpairs[i] &
		design$amount == results.by.dilution$amount[i] &
		design$cell %in% two.cells
	)
	if (length(which.samples) != 4) NA else {
		these.counts <- do.call(cbind, counts.list[which.samples])
		these.counts <- these.counts[rowSums(these.counts) > 0,]
		if (sum(! apply(these.counts == 0, 1, any)) == 0) NA else { # DESeq requires some all-nonzero rows
			colnames(these.counts) <- design$library[which.samples]
			result <- results(DESeq(DESeqDataSetFromMatrix(these.counts, colData = data.frame(cell = design$cell[which.samples]), design = ~ cell), fitType = "local"))
			total <- sum(! is.na(result$padj) & result$padj < 0.01)
			total
		}
	}
})))
plot.sig.promoters <- ggplot(subset(results.by.dilution, ! is.na(sig.promoters)), aes(readpairs / 1E6, sig.promoters, color = amount, group = amount)) +
	geom_line() +
	scale_x_continuous(name = "read pairs sequenced per library", label = unit_format(unit = "M", sep = "")) +
	scale_y_continuous(name = "differentially methylated promoters (adjusted p < 0.01)", label = comma, limit = c(0, NA)) +
	theme(legend.position = "n")
ggsave("subsample_sig_promoters.pdf", plot.sig.promoters, width = 4, height = 4)

# cluster silhouette
# uses results.by.dilution to get whole matrices but puts results in results.by.library
results.by.library$silhouette <- NA
for (results in mclapply(1:nrow(results.by.dilution), function(i) {
	which.samples <- which(
		design$readpairs == results.by.dilution$readpairs[i] &
		design$amount == results.by.dilution$amount[i] &
		design$cell %in% two.cells
	)
	if (length(which.samples) != 4) return(NULL) else {
		these.log10rpmpm <- do.call(cbind, log10rpmpm.list[which.samples])
		return(list(which.samples = which.samples, silhouettes = silhouette(as.integer(factor(design$cell[which.samples])), as.dist(1 - cor(these.log10rpmpm, use = "p")))[,"sil_width"]))
	}
})) if (! is.null(results)) results.by.library$silhouette[results$which.samples] <- results$silhouettes
plot.silhouette <- ggplot(subset(results.by.library, ! is.na(silhouette)), aes(readpairs / 1E6, silhouette, color = cell, group = library)) +
	geom_line() +
	facet_grid(amount ~ .) +
	scale_x_continuous(name = "read pairs sequenced per library", label = unit_format(unit = "M", sep = "")) +
	scale_y_continuous(name = "silhouette metric of cluster distance", limits = 0:1) +
	theme(legend.position = "n")
ggsave("subsample_silhouette.pdf", plot.silhouette, width = 3, height = 6)


save.image("analyze_subsamples.RData")
```
