## Supplemental Code for "Cost-effective DNA methylation profiling by FML-seq": subsample_promoter_hit.sh

```
#! /bin/bash

# subsample read pairs from the CRAM files to specifically targeted sequencing depths and redo the promoter-hit counting
# subsampling the final counts would not be the same because removing a read pair removes two connected hits
# samtools view --subsample does subsample pairs rather than individual reads, as desired
# target total sequenced read counts, not aligned counts or whatever
# get original read counts from bwa_stats.tsv

amount=60ng
increment=1000000

increments=0
echo 'found libraries and pair counts:'
while read library n_reads; do
	n_pairs=$(("$n_reads" / 2))
	libraries+=("$library")
	pairs+=($n_pairs) # divide n reads by 2 to get n pairs
	mkdir -p "subsample/$library"
	echo "$library" $n_pairs
	n_increments=$(($n_pairs / $increment))
	(($n_increments > $increments)) && increments=$n_increments
done < <(tail -n +2 bwa_stats.tsv | cut -f 1-2 | grep "$amount")
echo -e "\nsubsampling $increments increments of $increment pairs"

for i in $(seq $increments); do
	targets+=($(($increment * $i)))
done

parallel '
	proportion=$(bc <<< "scale=8; {3} / {2}")
	if [ $(bc <<< "$proportion < 1") -eq 1 ]; then
		samtools view -s $proportion -uq 10 -F 0x4 -F 0x100 -F 0x200 -F 0x800 cram/{1}.cram |
			~/methyl/count_methyl_hits.py -qw 0 <(zstdcat ~/genome/hs1/motif/CGNR_hs1.bed.zst) 2>/dev/null |
			~/methyl/region_counts.py -qz ~/genome/hs1/hs1.chrom.sizes ~/genome/hs1/ENCODE_hs1_cCREs_promoter-like.150-350bp.bed > subsample/{1}/{3}.tsv
		echo "subsampled {1} to {3} pairs from {2} ($proportion)"
	fi
' ::: "${libraries[@]}" :::+ "${pairs[@]}" ::: "${targets[@]}"
```
